# Maternal γδ T cells shape offspring pulmonary type-2 immunity in a microbiota-dependent manner

**DOI:** 10.1101/2021.08.13.456265

**Authors:** Pedro H. Papotto, Bahtiyar Yilmaz, Gonçalo Pimenta, Sofia Mensurado, Carolina Cunha, Gina Fiala, Daniel Gomes da Costa, Natacha Gonçalves-Sousa, Brian H. K. Chan, Birte Blankenhaus, Tânia Carvalho, Andrew J. Macpherson, Judith E. Allen, Bruno Silva-Santos

**Affiliations:** Instituto de Medicina Molecular João Lobo Antunes, Faculdade de Medicina, Universidade de Lisboa, Lisbon, Portugal; Lydia Becker Institute for Immunology & Inflammation, Faculty of Biology, Medicine & Health, Manchester Academic Health Science Centre, University of Manchester, Manchester, United Kingdom; Maurice Müller Laboratories, Department for Biomedical Research, University of Bern, Bern, Switzerland; Department of Visceral Surgery and Medicine, Bern University Hospital, University of Bern, Bern, Switzerland; Wellcome Centre for Cell-Matrix Research, Faculty of Biology, Medicine & Health, Manchester Academic Health Science Centre, University of Manchester, Manchester, United Kingdom

## Abstract

Immune system development is greatly influenced by vertically transferred cues. However, beyond antibody-producing B cells, little is known about the role of other cell subsets of the maternal immune system in regulating offspring immunity. We reasoned γδ T cells to be attractive candidates based on their tissue distribution pattern: abundant in the skin, mammary glands and female reproductive tract. Here we found that mice born from γδ T cell-deficient (TCRδ^-/-^) dams display, early after birth, a pulmonary milieu selectively enriched in type-2 cytokines such as IL-33, IL-4, IL-5, and IL-13, and type-2-polarized immune cells, when compared to the progeny of γδ T cell-sufficient dams. In addition, upon helminth infection, mice born from TCRδ^-/-^ dams sustained an increased type-2 inflammatory response. Critically, this was independent of the genotype of the pups. Despite similar levels of circulating antibodies in mothers and progeny, the intestinal microbiota in the offspring of TCRδ^-/-^ and TCRδ^+/-^ dams harbored distinct bacterial communities acquired during birth and fostering. These differences were accompanied by changes in the intestinal short-chain fatty acids (SCFA) profile. Importantly, antibiotic treatment abrogated the differences observed in the pulmonary milieu, and exogenous SCFA supplementation suppressed first-breath- and infection-induced inflammation. In summary, maternal γδ T cells control the establishment of a neonatal gut–lung axis by conditioning the postnatal microbial colonization of the offspring and bacterial-derived metabolite availability; ultimately impacting on the development of pulmonary type-2 immunity in the offspring.

## Introduction

The development of the immune system is particularly sensitive to maternally derived factors. Maternal transfer of bacterial species during pregnancy and pre-weaning is responsible for determining the offspring’s microbiota profile, which in turn influences immune responses later in life *(Al Nabhani et al., 2019; Tanaka and Nakayama, 2017; Wampach et al., 2018)*, even if strong causal relationships are yet to be demonstrated. This notwithstanding, vertical transfer of bacterial products via antibodies during pregnancy was shown to modulate the maturation of the intestinal immune system in the offspring *(Gomez de Aguero et al., 2016)*.

Maternally transferred antibodies were also recently shown to control the homeostatic number of colonic regulatory T (Treg) cells across generations *(Ramanan et al., 2020)*. Furthermore, maternal exposure to a high fiber diet during pregnancy leads to decreased airway allergic disease severity in their progeny, through the transfer of short-chain fatty acids (SCFA), a major class of dietary metabolites *(Thorburn et al., 2015)*. Interestingly, airway immune disorders seem to be tightly connected with early life exposure to environmental challenges *(Renz and Skevaki, 2021)* likely due to type-2 immune activation and tissue remodeling characterizing the perinatal phase of lung development *(de Kleer et al., 2016; Saluzzo et al., 2017)*. Apart from exciting biological consequences, this maternal crosstalk has profound implications on the design of experiments using genetically engineered mice, namely the choice of controls to generate accurate and reproducible data. Even though littermate controls are considered the gold standard *(Stappenbeck and Virgin, 2016)*, it is clear that alternative approaches, such as co-housing and especially genetic background matching (to so-called “wild type” mice, often obtained directly from commercial vendors), are still widely used in biomedical research. However, given that many external cues are vertically transferred during early life and have long-lasting consequences *(Gomez de Aguero et al., 2016; Kimura et al., 2020; Ramanan et al., 2020; Thorburn et al., 2015)*, controlling for environmental factors without using littermate controls is nearly impossible. On a positive note, the problems evidenced by the use of sub-optimal controls raise interesting questions on how parental-derived factors affect the offspring *(Macpherson et al., 2017; Perez and Lehner, 2019)*. In particular, even though the maternal influence on progeny immune system is evident, the role of individual components of the maternal immune system (beyond antibodies) in regulating offspring immunity remains largely unexplored.

γδ T cells are a population of innate-like lymphocytes known to reside in different tissues that compose the maternal-newborn interface, such as the female reproductive tract *(Itohara et al., 1990; Monin et al., 2020; Pinget et al., 2016)*, the skin *(Koning et al., 1987; Nielsen et al., 2017)* and mammary gland *(Reardon et al., 1990)*. Murine γδ T cells were shown to control antibody production both under steady state conditions *(Fujihashi et al., 1996)* and upon challenge *(Rezende et al., 2018)*. Also, γδ T cells are known to participate in an intricate crosstalk with barrier tissue microbiota *(Khairallah et al., 2018; Papotto et al., 2021)*, notably being able to exert significant selective pressure on bacterial communities, as recently documented for the oral microbiome *(Krishnan et al., 2018; Wilharm et al., 2019)*. Hence, it is conceivable that maternal γδ T cells affect offspring immune system development through a variety of mechanisms, but these have yet to be investigated. Here we show that pups born from TCRδ^-/-^ (compared to TCRδ^+/-^) dams present a perinatal pulmonary environment strongly biased towards type-2 immunity, both in the steady state and upon helminth infection, irrespectively of their own genotype. Based on cross-fostering and cesarean section (C-Sec) delivery experiments, as well as antibiotic treatment during gestation and nursing, we propose a critical role for the microbiota in this process. Consistently, pups born from TCRδ^-/-^ dams display distinct intestinal microbial communities and reduced intestinal SCFA. Importantly, exogenous administration of SCFA to mice born from TCRδ^-/-^ dams decreases perinatal type-2 inflammation and suppresses the immune response to helminth infection. Overall, these data suggest that maternal γδ T cells regulate postnatal microbial colonization and microbial derived metabolite availability in the offspring, ultimately impacting on their pulmonary immune system development.

## Results

### Absence of maternal γδ T cells exacerbates perinatal type-2 pulmonary inflammation in offspring

The importance of pulmonary γδ T cells in the establishment of lung type-2 immune responses has been highlighted by various groups *(Ajendra et al., 2020; Sutherland et al., 2014; Van Dyken et al., 2014)*. While trying to understand the mechanisms underlying the γδ T cell – type-2 immunity crosstalk, we observed that mice from our TCRδ^-/-^ colony possessed decreased frequencies and numbers of pulmonary type-2 innate lymphoid cells (ILC2), when compared to age- and sex-matched adult C57BL6/J mice **(Figures S1A and S1B)**. However, and intriguingly, when we compared TCRδ^+/+^ and TCRδ^-/-^ littermates from heterozygous breedings (TCRδ^+/-^ × TCRδ^+/-^), no differences were observed in lung ILC2 **(Figure S1C and S1D)**. As our TCRδ^+/-^ colony is derived from the TCRδ^-/-^ colony and both experience the same environmental factors, we hypothesized that absence of γδ T cells in the dams could be responsible for altered lung ILC2. To test this hypothesis, we established reciprocal breeding colonies by crossing TCRδ^-/-^ females with TCRδ^+/-^ males and vice-versa; these breeding strategies give rise to both TCRδ-/- and TCRδ+/- progeny allowing us to exclude gene-intrinsic effects in the offspring **(Figure 1A)**. Importantly, we chose to analyze the lungs of pups born from γδ-deficient and -sufficient dams around post-natal day (PN) 15, as the main events leading to the maturation of the pulmonary immune system peak during this time as a consequence of the first-breath-induced inflammatory reaction *(de Kleer et al., 2016; Saluzzo et al., 2017)*. In accordance with our hypothesis, pups born from TCRδ^-/-^ dams exhibited a decrease in lung ILC2, when compared to pups born from TCRδ^+/-^ dams **(Figures S1E and S1F)**. Interestingly, the offspring of TCRδ^-/-^ dams displayed augmented perinatal pulmonary type-2 inflammation, including increases in IL-13^+^ ILC2 **(Figures 1B–C)**, and IL-5^+^ mast cells **(Figures S1I–K)**. Accordingly, we also observed increased tissue levels of the type-2-inducing alarmin IL-33, and the type-2 cytokines IL-4 and IL-5, in the lungs of the progeny of TCRδ^-/-^ dams, when compared with pups born from their TCRδ^+/-^ counterparts **(Figure 1E)**. In addition, consistent with findings in type-2 asthma models (Walter et al., 2001), IL-12p40, but not IL-12p70 and IL-23, was also increased in the lungs of the offspring from TCRδ^-/-^ dams **(Figures 1E and S1N)**. Moreover, interstitial macrophages from the offspring of TCRδ^-/-^ dams exhibited increased expression of type-2-regulated genes, such as *Il13ra, I1rl1*, and *Arg1* **(Figure 1F and 1G)**. Critically, when the same pups were stratified according to their own genotype, no differences were observed between TCRδ^-/-^ and TCRδ^+/-^ pups regarding lung ILC2 **(Figures 1H, 1I, S1G and S1H)**, mast cells **(Figures S1L and S1M)**, lung cytokine levels **(Figure 1J)**, or gene expression on interstitial macrophages (Figure 1K). Thus, the observed increase in first-breath-induced type-2 immune responses in pups born from TCRδ^-/-^ dams depends solely on the maternal genotype. Also, these results stress the importance of using littermates to control for vertically-transferred factors. Despite being considered “gold standard”, this practice it is still not common in biomedical research; in particular, in the field of γδ T cells and infectious diseases, only about 15% of the published studies since the development of TCRδ^-/-^ mice (1993) have used littermate controls over the years **(Figure S2A–B)**, including a notable absence from top-ranked journals **(Figure S2C)**.

**Fig. 1.**
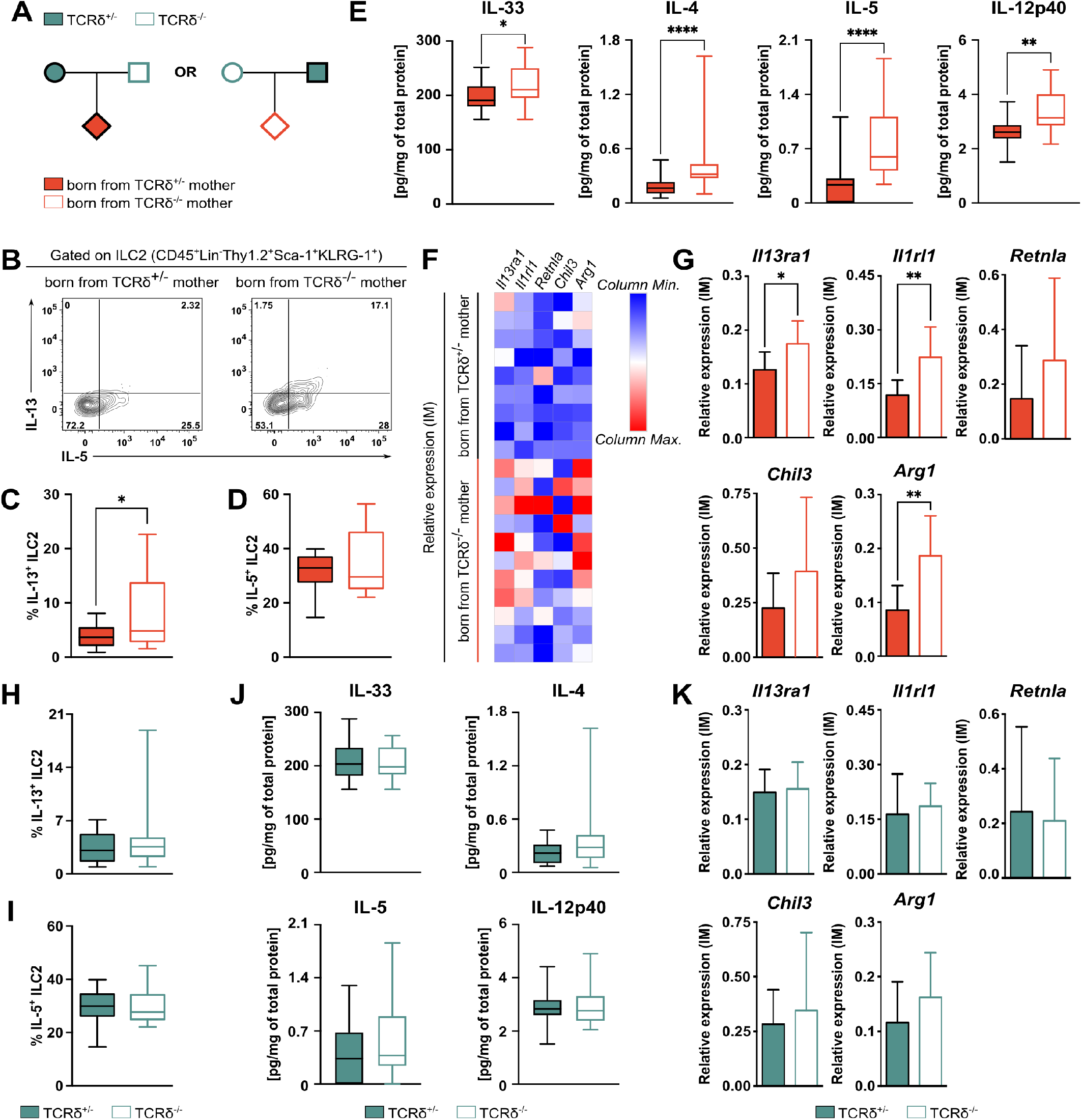
Absence of maternal γδ T cells exacerbates first-breath-induced type-2 inflammation in the offspring. **(A)** Breeding strategy employed to evaluate the gene-extrinsic and gene-intrinsic roles of γδ T cell depletion on the formation of neonatal pulmonary immune system. TCRδ^+/-^ dams were crossed with TCRδ^-/-^ males, and vice-versa; the pups were kept with the parents and analyzed at PN16±2. **(B)** Flow cytometry analysis of intracellular IL-5 and IL-13 in ILC2s (defined as Lin-Thy1.2^+^Sca-1^+^KLRG-1^+^ cells) isolated from the lungs of pups generated in the breedings described in (A) and stimulated in vitro for 3h in the presence of PMA, Ionomycin and Brefeldin A. Frequencies of **(C)** IL-13^+^ and **(D)** IL-5^+^ cells within the ILC2 population in the lungs of pups depicted in (A), grouped by maternal genotype. **(E)** Concentration of cytokines within the whole lung homogenate normalized by total protein in the tissue of pups from the breedings described in (A), grouped by maternal genotype. **(F)** Heatmap depicting mRNA expression (relative to *Hprt* and *Actb*) of selected genes by qPCR in sorted interstitial macrophages (IM; defined as CD45^+^CD64^+^CD24^-^CD11c^-^CD11b^+^ cells) from the pups generated in the breedings described in (A), grouped by maternal genotype. **(G)** mRNA expression (relative to *Hprt* and *Actb*) of selected genes by qPCR in sorted IM from the pups generated in the breedings described in (A), grouped by maternal genotype. Frequencies of **(H)** IL-13^+^ and **(I)** IL-5^+^ cells within the ILC2 population in the lungs of pups depicted in (A), grouped by offspring genotype. **(J)** Concentration of cytokines within the whole lung homogenate normalized by total protein in the tissue of pups from the breedings described in (A), grouped by offspring genotype. **(K)** mRNA expression of selected genes by qPCR in sorted IM from the pups generated in the breedings described in (A), grouped by offspring genotype. **(B-D; H-I)** Data pooled from three independent litters per maternal genotype. n= 16-17 mice per group. **(E-J)** Data pooled from at least three independent litters per genotype; n= 18-23 mice per group. **(G, K)** n= 9-11 mice per group **(C-E; H-J)** Box-and-whisker plots display first and third quartiles, and the median; whiskers are from each quartile to the minimum or maximum. **(G, K)** Error bars represent Mean ±SD. Normality of the samples was assessed with D’Agostino Pearson normality test; statistical analysis was then performed using Student’s t test or Mann-Whitney test. * *p* < 0.05; ** *p* < 0.01; **** *p* < 0.0001

Of note, despite the differences observed in pulmonary type-2 immunity, pups born from TCRδ^-/-^ and TCRδ^+/-^ dams presented a similar distribution of lung myeloid populations **(Figure S3A)**, T and B lymphocytes **(Figure S3B)**, and regulatory T cells **(Figure S3C)**. Moreover, the expression of epithelial genes controlling surfactant and mucin production, and barrier integrity was similar between pups from both breeding strategies **(Figure S3D)**. Finally, the differences between the progeny of TCRδ^-/-^ and TCRδ^+/-^ dams seemed restricted to the lung, as we found similar frequencies of ILC2 precursors and other lymphoid and myeloid progenitors in the bone marrow **(Figure S4A)**, and of innate lymphoid cells in the small intestine lamina propria **(Figure S4B)**. These data indicate that maternal γδ T cells “vertically” (and selectively) regulate first-breath-induced pulmonary type-2 inflammation in the offspring.

### Microbiota transfer in early life mediates differences in the pulmonary immune system of the offspring from TCRδ^+/-^ and TCRδ^-/-^ dams

Transfer of maternally-derived factors during pregnancy has been shown to regulate the development of the intestinal immune system *(Gomez de Aguero et al., 2016; Ramanan et al., 2020)* and suppress airway inflammation in the offspring *(Thorburn et al., 2015)*. Our results indicated that a similar process could also fine tune first-breath-induced inflammation. Surprisingly, cross-fostering pups born from TCRδ^-/-^ dams with TCRδ^+/-^ surrogates, and vice-versa **(Figure 2A)**, abrogated the differences observed in the pulmonary milieu between their respective progeny **(Figure 2B, compared to Figure 1E)**. This suggested that the vertical conditioning of offspring lung immunity was not taking place exclusively either during pregnancy or fostering. In order to further exclude a role for maternally-transferred factors during gestation and lactation, pups born via C-Sec from either TCRδ^-/-^ or TCRδ^+/-^ dams were co-fostered with pups born naturally (NB) by a mother of the same genotype **(Figure 2C)**. Consistently, whereas NB pups from TCRδ^-/-^ dams retained increased lung levels of IL-33, and IL-13-producing pulmonary ILC2, C-Sec pups born from TCRδ^-/-^ dams displayed no signs of increased perinatal type-2 inflammation in the lungs in comparison to control pups **(Figures 2D–G)**. Altogether, these data reveal the period around delivery to be a critical time window for the transfer of maternally-derived factors, which in turn regulates perinatal lung inflammation in the offspring.

**Fig. 2.**
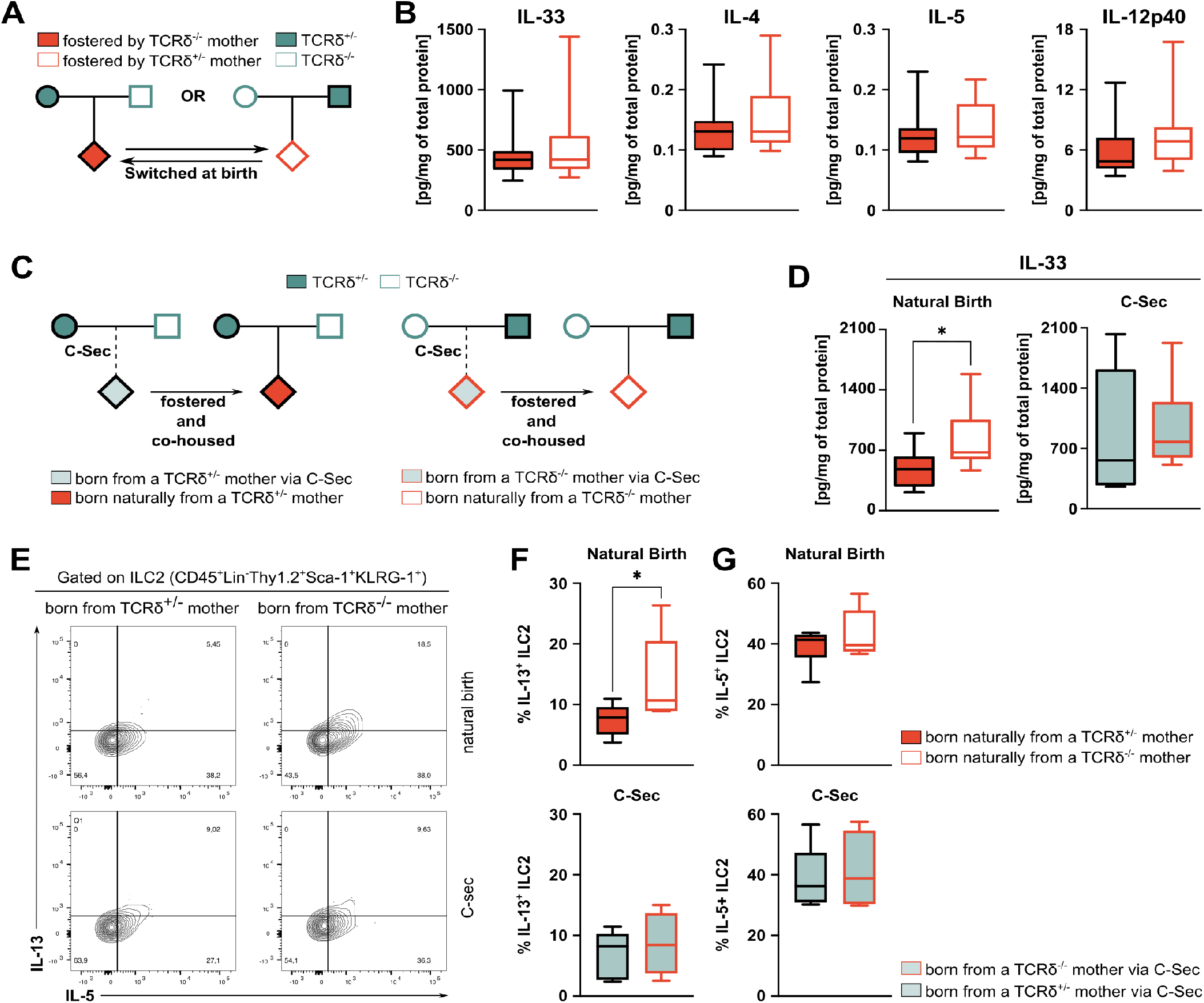
Maternal regulation of first-breath-induced inflammation in the offspring occurs during delivery and fostering. **(A)** Breeding strategy employed to evaluate the role of fostering on perinatal lung inflammation. TCRδ^+/-^ dams were crossed with TCRδ^-/-^ males, and vice-versa; pups were then swapped between mothers one day after birth and kept with the dams until weaning, and analyzed at PN16±2. **(B)** Concentration of cytokines within the whole lung homogenate of pups from the breedings described in (A), grouped by maternal genotype; concentration was normalized to the total protein in the tissue. **(C)** Breeding strategy employed to evaluate the role of delivery mode on perinatal lung inflammation. TCRδ^+/-^ dams were crossed with TCRδ^-/-^ males, and vice-versa; part of the pregnant females was subjected to C-Sec surgery, and their pups were then fostered by a mother with the same genotype that had undergone normal delivery one day before. C-Sec pups were co-housed with naturally delivered neonates, and analyzed at PN16±2. **(D-G)** Concentration of cytokines within the whole lung homogenate of pups from the breedings described in (C), grouped by maternal genotype; concentration was normalized to the total protein in the tissue. **(H)** Flow cytometry analysis of intracellular IL-5 and IL-13 in ILC2s (defined as Lin^-^Thy1.2^+^Sca-1^+^KLRG-1^+^ cells) isolated from the lungs of pups generated in the breedings described in (C) and stimulated in vitro for 3h in the presence of PMA, Ionomycin and Brefeldin A. Frequencies of **(I)** IL-13^+^ and **(J)** IL-5^+^ cells within the ILC2 population in the lungs of pups depicted in (C), grouped by maternal genotype and mode of delivery. **(B)** Data pooled from three independent litter swaps per maternal genotype; n= 18-21 mice per group. **(D)** Data pooled from two independent litters per maternal genotype; n= 9-11 mice per group. **(E-G)** Data from one litter per maternal genotype; n= 4-6 mice per group. **(B; D; F-G)** Box-and-whisker plots display first and third quartiles, and the median; whiskers are from each quartile to the minimum or maximum. Normality of the samples was assessed with D’Agostino Pearson normality test; statistical analysis was then performed using Student’s t test or Mann-Whitney test. * *p* < 0.05

Maternal antibodies have been shown to prevent the development of asthma in the offspring through the control of aberrant type-2 immune responses *(Ohsaki et al., 2018)*, and γδ T cells can regulate natural and adaptive antibody production *(Fujihashi et al., 1996; Rezende et al., 2018)*. However, it seemed unlikely that maternal transfer of antibodies during delivery would be playing a role in our case. Indeed, the circulating levels of total IgG and IgG subclasses were similar between both TCRδ^-/-^ and TCRδ^+/-^ dams **(Figures 3A and 3B)** and their respective progeny **(Figures 3A and 3C)**. To further confirm that maternal antibodies were not involved in the regulation of first-breath-induced inflammation, we established reciprocal breedings by crossing antibody-deficient JHT females with antibody-sufficient males, and vice-versa **(Figure 3D)**. As expected, the offspring of both JHT dams and their WT counterparts presented similar tissue levels of type-2 cytokines IL-33, IL-4, and IL-5 in their lungs, but a slight increase in IL-12p40 **(Figure 3E)**.

**Fig. 3.**
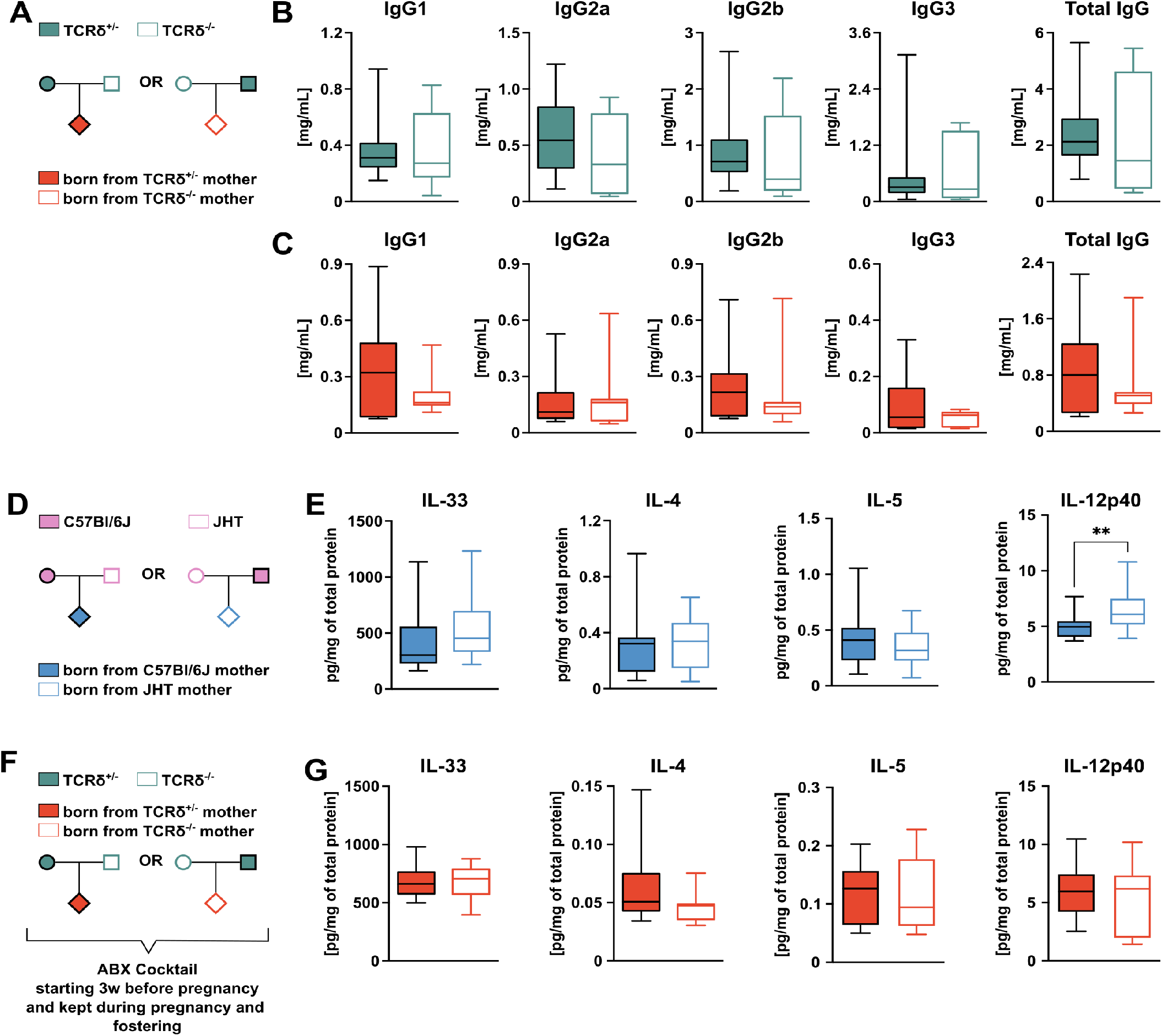
Maternal regulation of first-breath-induced inflammation in the offspring is microbiota-dependent but antibody-independent. **(A)** Breeding strategy employed to evaluate the gene-extrinsic and gene-intrinsic roles of γδ T cell depletion on the formation of neonatal pulmonary immune system. TCRδ^+/-^ dams were crossed with TCRδ^-/-^ males, and vice-versa; the pups were kept with the parents until weaning, and analyzed at PN16±2. **(B)** Concentration of total IgG and its subclasses in the serum of TCRδ^+/-^ and TCRδ^-/-^ dams. **(C)** Concentration of total IgG and its subclasses in the serum of pups born from TCRδ^+/-^ and TCRδ^-/-^ dams. **(D)** Breeding strategy employed to evaluate if maternal transfer of antibodies impact on perinatal lung inflammation. JHT dams were crossed with C57Bl6/J males, and vice-versa; the pups were kept with the parents until weaning, and analyzed at PN16±2. **(E)** Concentration of cytokines within the whole lung homogenate normalized by total protein in the tissue of pups from the breedings described in (A), grouped by maternal genotype. **(F)** Breeding strategy employed to evaluate the role of microbiota in mediating maternal γδ T cell conditioning of the pulmonary immune system of the offspring. TCRδ^+/-^ and TCRδ^-/-^ males and females were treated for 3 weeks prior breeding setup with a broad-spectrum antibiotic cocktail containing streptomycin, ampycilin, collistin and vancomycin; TCRδ^+/-^ females were then crossed with TCRδ^-/-^ males, and vice-versa. Antibiotic treatment was kept during pregnancy and fostering. The pups were kept with the parents, and analyzed at PN16±2. **(G)** Concentration of cytokines within the whole lung homogenate normalized by total protein in the tissue of pups from the breedings described in (F), grouped by maternal genotype. **(B)** n= 7-14 mice per group. (C) Data pooled from three independent litters per maternal genotype; n= 19-21 mice per group. **(E)** Data pooled from three independent litters per maternal genotype; n= 18-22 mice per group. **(G)** Data pooled from three independent litters per maternal genotype, n= 17-18 mice per group. **(B-C; E; G)** Box-and-whisker plots display first and third quartiles, and the median; whiskers are from each quartile to the minimum or maximum. Normality of the samples was assessed with D’Agostino Pearson normality test; statistical analysis was then performed using Student’s t test or Mann-Whitney test. ** *p* < 0.01

The period comprising delivery and weaning is critical for the formation of neonatal microbiota. Importantly, vertical transfer of microbiota *(Wampach et al., 2018)* or its products *(Gomez de Aguero et al., 2016)* have also been implicated in the modulation of offspring immunity. Hence, we treated parents and progeny from our reciprocal breeding strategies with a wide spectrum antibiotics (ABX) cocktail, starting from before pregnancy through pregnancy and fostering **(Figure 3F)**. Interestingly, in contrast with the control setting **(Figure 1E)**, pups born from ABX-treated TCRδ^-/-^ and TCRδ^+/-^ dams presented similar tissue levels of cytokines in their lungs **(Figure 3G)**, suggesting that the enhanced type 2 pulmonary responses observed in pups born to TCRδ^-/-^ dams resulted from vertical transfer of microbiota.

### TCRδ^-/-^ dams co-housed with TCRδ^+/-^ males retain differences in cutaneous bacterial communities

Co-housing mice with a different starting microbiota highly homogenizes the microbiota between them *(Caruso et al., 2019)*. Thus, for our breeding setup, we expected to observe no differences in the microbiota of TCRδ^-/-^ and TCRδ^+/-^ dams, upon co-housing with a male from the opposing genotype. In fact, before establishment of the reciprocal breedings, TCRδ^-/-^ and TCRδ^+/-^ females displayed significant differences in fecal bacterial diversity and richness **(Figures S5A and S5B)**. However, after breeding establishment, beta diversity analysis of the fecal microbiota showed a significant change in bacterial diversity of TCRδ^-/-^ and TCRδ^+/-^ females before and after co-housing **(Figures S5C and S5D)**; no changes in bacterial richness were observed (Figures S5E and S5F). Accordingly, after establishment of the breedings, TCRδ^-/-^ and TCRδ^+/-^ dams displayed similar bacterial diversity and richness in their fecal **(Figures S6A and S6B)**, vaginal **(Figures S6C and S6D)** and cutaneous **(Figures S6E and S6F)** microbiota. This notwithstanding, comparison of the relative abundances of specific skin microbial communities showed a significant increase of Bacteroidetes phyla and *Oscillospira* genus in TCRδ^+/-^ dams **(Figures S6G and S6H)**, and a minor increase of *Parvimonas* and *Tannerella* genera in TCRδ^+/-^ dams **(Figures S6G)**. Hence, even though the co-housing process largely homogenizes the microbiota, taxa-specific differences in the skin microbiota are retained between TCRδ^+/-^ and TCRδ^-/-^ dams.

### Decreased availability of microbial-derived SCFA in the progeny of TCRδ^-/-^ dams causes exacerbated pulmonary type-2 inflammation

The data from the C-sec experiment, together with the differences observed in the cutaneous microbiota of γδ T cell-sufficient and -deficient dams, led us to hypothesize that their offspring are subjected to a distinct neonatal microbial colonization. Bacterial composition of the cecal contents from the offspring of TCRδ^-/-^ and TCRδ^+/-^ dams evidenced similar bacterial diversity and richness independently on maternal and progeny genotype **(Figure S7A and S7B)**. However, pups born from TCRδ^+/-^ dams displayed increased relative abundances of Bacteroidales order, Ruminococcaceae and S24-7 families, and *Clostridium* and *Adlercreutzia* genera, while in contrast the *Clostridium* genus was relatively more abundant in pups born from TCRδ^-/-^ dams **(Figures 4A and S7C)**. Most importantly, to evaluate the relevance of these taxonomy changes, we used the PICRUSt algorithm to predict the functional metagenome of cecal bacterial communities found in the offspring of TCRδ^-/-^ and TCRδ^+/-^ dams *(Langille et al., 2013)*. Several metabolic pathways were predicted to be upregulated in the microbiota of pups born from TCRδ^+/-^ dams, most of which related to the fermentation of SCFA **(Figure 4B)**. Corroborating our in-silico predictions, analysis of cecum levels of SCFA by gas-chromatography–mass-spectrometry showed that the progeny of TCRδ^-/-^ and TCRδ^+/-^ dams possess distinct SCFA profiles **(Figure 4C)**, with pups born from TCRδ^+/-^ dams presenting increased levels of hexanoate and pentanoate and, although not statistically significant (*p* = 0.0506), a trend for increased levels of acetate; no differences were observed in the other SCFA measured **(Figures 4D and S7D)**. Of note, we did not observe differences in bacterial diversity and richness in the lung microbiota from the offspring of TCRδ^-/-^ and TCRδ^+/-^ dams **(Figures S7E and S7F)**. Overall, these results reveal a distinct intestinal bacterial colonization and availability of microbial-derived metabolites between pups born from TCRδ^-/-^ and TCRδ^+/-^ dams, suggesting the possibility of a gut-lung axis linking the bacterial-derived SCFA and the regulation of first-breath-induced inflammation.

**Fig. 4.**
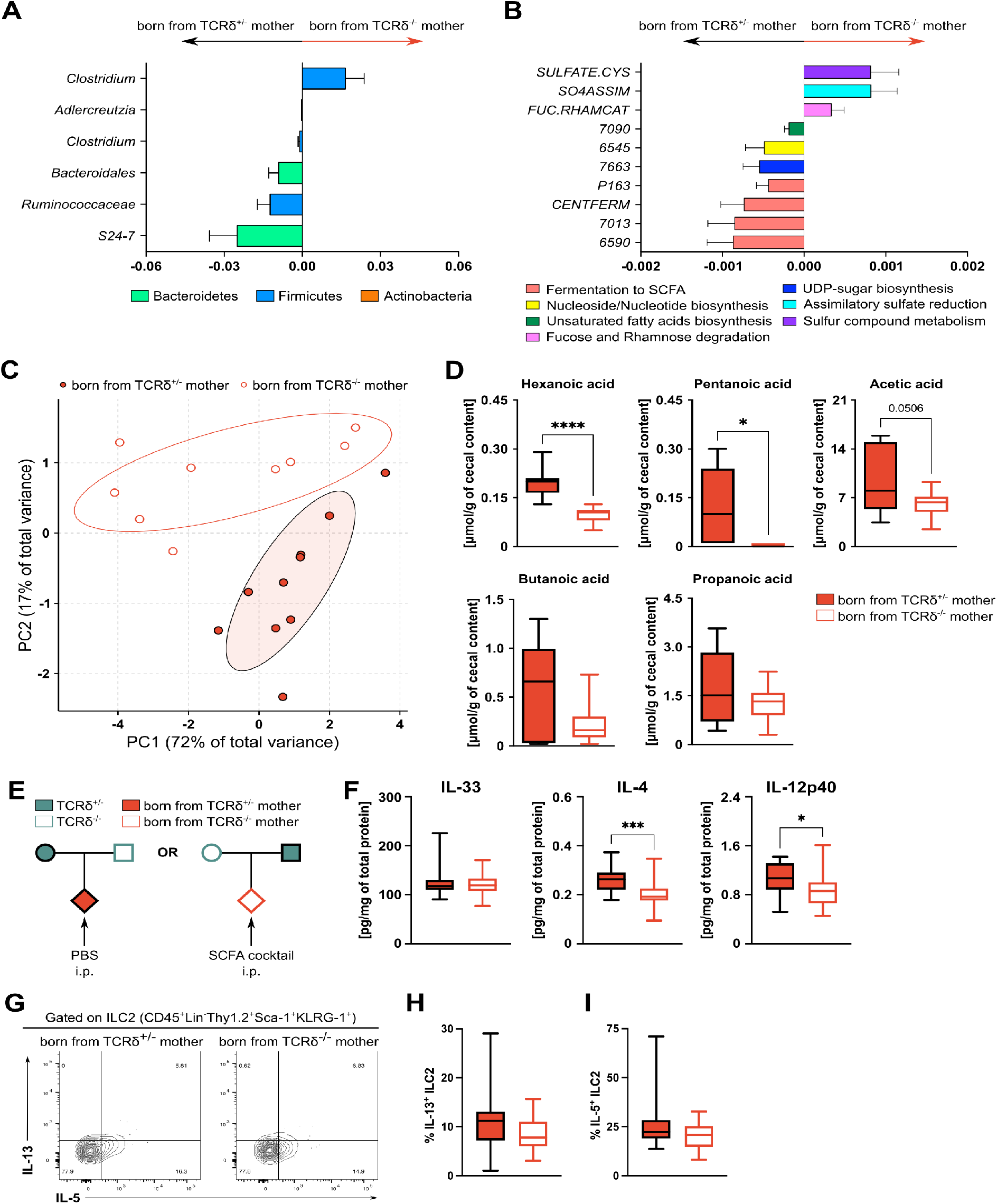
Microbiota-derived SCFA in the offspring suppress exacerbated perinatal type-2 inflammatory responses in the lung. **(A)** Cecum microbiota taxa associated with TCRδ^+/-^ and TCRδ^-/-^ maternal genotypes and plotted as relative abundance ratios. **(B)** PICRUSt analysis of the cecum microbiota of pups born from TCRδ^+/-^ and TCRδ-/- dams and plotted as differentially regulated pathways associated with maternal genotype. **(C)** Principal Component Analysis (PCA) plot depicting the clustering of pups born from TCRδ^+/-^ and TCRδ^-/-^ dams based on their cecal SCFA profile. **(D)** Concentration of SCFA present in the cecum of pups born from TCRδ^+/-^ and TCRδ^-/-^ dams normalized by cecal content. **(E)** Breeding strategy employed to evaluate the role of SCFA in regulating perinatal lung inflammation. TCRδ+/- dams were crossed with TCRδ^-/-^ males, and vice-versa; pups were kept with the parents and administered a SCFA cocktail (acetate, pentanoate and hexanoate) i.p. every three days, starting at PN7. Analysis of neonatal lung immune system was conducted at PN16±2. **(F)** Concentration of cytokines within the whole lung homogenate normalized by total protein in the tissue of pups from the breedings described in (E), grouped by maternal genotype. **(G)** Flow cytometry analysis of intracellular IL-5 and IL-13 in ILC2s (defined as Lin^-^Thy1.2^+^Sca-1^+^KLRG-1^+^ cells) isolated from the lungs of pups generated in the breedings described in (E) and stimulated in vitro for 3h in the presence of PMA, Ionomycin and Brefeldin A. Frequencies of **(H)** IL-13^+^ and **(I)** IL-5^+^ cells within the ILC2 population in the lungs of pups depicted in (E), grouped by maternal genotype. **(A-B)** Data pooled from two independent litters per maternal genotype; n= 10-14 mice per group. **(C-D)** Data from one litter per maternal genotype; n= 9-10 mice per group. (F-I) Data pooled from three independent litters per maternal genotype; n= 17-18 mice per group. **(A-B)** Error bars represent Mean ±SD; Statistical analyses for taxonomy and PICRUSt results were performed using the MaAsLin pipeline with Benjamini–Hochberg false-discovery rate (BH-FDR) (q value) q < 0.2 and p < 0.05 being considered significant **(D; F; H-I)** Box-and-whisker plots display first and third quartiles, and the median; whiskers are from each quartile to the minimum or maximum; Normality of the samples was assessed with D’Agostino Pearson normality test; statistical analysis was then performed using Student’s t test or Mann-Whitney test. * *p* < 0.05; *** *p* < 0.001; **** *p* < 0.0001

In order to test if exogenous SCFA supplementation would be able to suppress the increased type-2 inflammation observed in the lungs of pups born from γδ T cell-deficient dams, we administered a SCFA cocktail to the offspring of TCRδ^-/-^ dams **(Figure 4E)**. The SCFA cocktail contained pentanoate and hexanoate (the two metabolites found to be significantly different between the pups), together with acetate, which, although not quite statistically significant, is the most abundant SCFA in the cecum of newborn mice **(Figure 4D)**. Confirming our hypothesis, administration of the SCFA cocktail to pups born from TCRδ^-/-^ dams abrogated the differences observed in the tissue levels of IL-33, whilst significantly decreasing IL-4 and IL-12p40 concentrations **(Figure 4F)**. Moreover, the treatment also decreased ILC2 activation in the offspring of TCRδ^-/-^ dams, as evidenced by similar frequencies of IL-13^+^ and IL-5^+^ cells **(Figures 4G–I)**. Importantly, administration of the SCFA cocktail via drinking water of TCRδ^-/-^ dams was not sufficient to decrease the frequencies of IL-13- and IL-5-producing ILC2 in their progeny when compared to that of their TCRδ^+/-^ counterparts **(Figure S8A–D)**; lung levels of IL-33 were also still higher in pups born from TCRδ^-/-^ dams, but not IL-4, IL-5 and IL-12p40 **(Figure S8E)**. Altogether, these findings indicate that bacterial but not maternally derived SCFA are critical in regulating the perinatal type-2 lung inflammation during steady state conditions.

### SCFA levels during fostering control post-weaning response to helminth infection

The increase in first-breath-induced type-2 inflammation in the pups born from γδ T cell-deficient dams prompted us to ask whether these mice would respond differently to a type-2-promoting insult. In order to address this question, mice were infected at the early post-weaning period with the helminth *Nippostrongylus brasiliensis* **(Figure 5A)**, a parasite that has been widely used as a model to elicit strong type-2 immunity in the lungs. Mice born from TCRδ^-/-^ dams presented an increased inflammatory response to the infection when compared to the offspring of γδ T cell-sufficient dams, characterized by higher numbers of total leukocytes infiltrating the lung **(Figure 5B)**, and enhanced eosinophilia and neutrophilia **(Figures 5C and 5D)**. Surface expression of Programmed Cell Death-Ligand 2 (PD-L2) in macrophages and dendritic cells (DCs) has been reported to occur under strong type-2 conditions *(Gundra et al., 2014; Selenko-Gebauer et al., 2003)*. Accordingly, we observed increased numbers of PD-L2-expressing macrophages **(Figures 5E and 5F)** and DCs **(Figures 5G and 5H)** in mice born from TCRδ-/- dams, when compared to controls. In line with these results, we also found an increase in IL-13^+^ and IL-17A^+^ (but not IFN-γ^+^) CD4^+^ T cells **(Figures 5I and 5J)** in the progeny of TCRδ^-/-^ dams. When mice were categorized according to their own genotype we observed no differences between TCRδ^-/-^ and TCRδ^+/-^ animals in any of the aforementioned parameters **(Figures S9A–F)**. Taken together, we have found that the offspring of TCRδ^-/-^ dams display stronger type-2 immune responses, both in steady-state and upon an infectious challenge, irrespective of their own genotype.

**Fig. 5.**
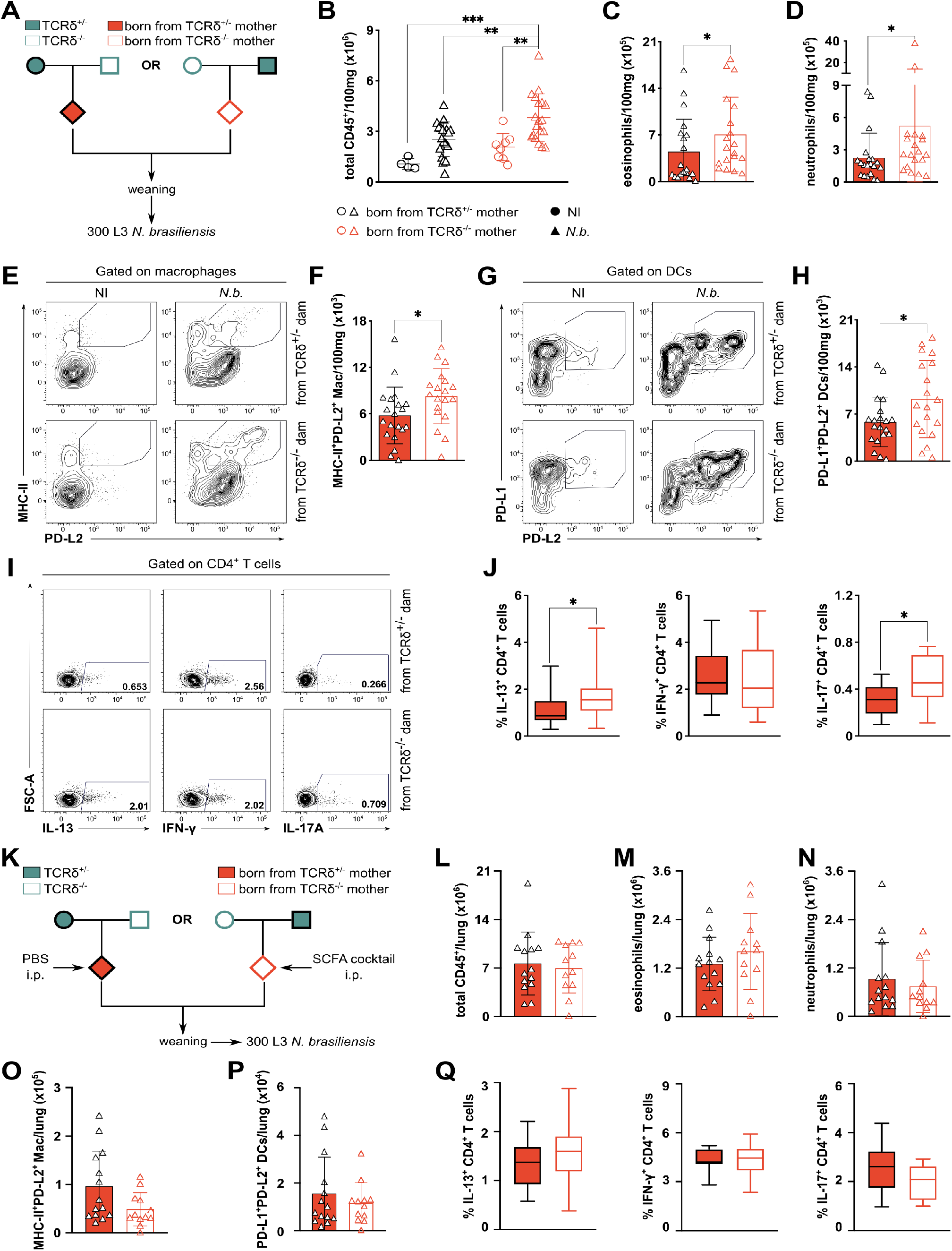
SCFA control exacerbated response to *N. brasiliensis* infection by the offspring of γδ T cell-deficient dams. **(A)** Breeding strategy employed to evaluate the gene-extrinsic and gene-intrinsic roles of γδ T cell depletion on pulmonary immunity. TCRδ^+/-^ dams were crossed with TCRδ^-/-^ males, and vice-versa; the pups were kept with the parents until weaning, to be later divided by sex. Mice were weaned at post-natal day (PN) 21 and then infected subcutaneously with 300 L3 from *N. brasiliensis* and analyzed at day 6 p.i. Numbers (normalized to 100mg of lung tissue) of **(B)** total leukocytes, **(C)** eosinophils, and **(D)** neutrophils in mice described in (A). **(E)** Flow cytometry analysis of surface MHC-II and PD-L2 expression in lung macrophages (defined as CD45^+^CD64^+^CD24^-^ live cells) isolated from the lungs of *N. brasiliensis*-infected mice from the breedings described in (A). **(F)** Numbers (normalized to 100mg of lung tissue) of MHC-II^+^PD-L2^+^ macrophages in the lungs of *N. brasiliensis*-infected mice from the breedings described in (A). **(G)** Flow cytometry analysis of surface PD-L1 and PD-L2 expression in lung dendritic cells (DCs; defined as CD45^+^CD64^-^CD24^+^CD11c^+^MHC-II^+^ live cells) isolated from the lungs of *N. brasiliensis*-infected mice from the breedings described in (A). **(H)** Numbers (normalized to 100mg of lung tissue) of PD-L1^+^PD-L2^+^ DCs in the lungs of N. brasiliensis-infected mice from the breedings described in (A). Flow cytometry analysis of IL-13^+^, IFN-γ^+^ and IL-17A^+^ CD4^+^ **(I)** and their frequencies **(J)** in the lungs of *N. brasiliensis*-infected mice from the breedings described in (A). **(K)** Breeding strategy employed to evaluate the role of SCFA on regulation of offspring pulmonary immunity. TCRδ^+/-^ dams were crossed with TCRδ^-/-^ males, and vice-versa; pups were kept with the parents and administered a SCFA cocktail (acetate, pentanoate and hexanoate) i.p. every three days, starting at PN14. Mice were weaned, and divided by sex at PN21 and then infected subcutaneously with 300 L3 from *N. brasiliensis*. Analysis was conducted at day 6 p.i. Numbers (normalized to 100mg of lung tissue) of **(L)** total leukocytes, **(M)** eosinophils, and **(N)** neutrophils in mice described in (K). **(O)** Numbers (normalized to 100mg of lung tissue) of MHC-II^+^PD-L2^+^ macrophages (defined as CD45^+^CD64^+^CD24^-^ live cells) in the lungs of *N. brasiliensis*-infected mice from the breedings described in (K). **(P)** Numbers (normalized to 100mg of lung tissue) of PD-L1^+^PD-L2^+^ DCs (defined as CD45^+^CD64^-^CD24^+^CD11c^+^MHC-II^+^ live cells) in the lungs of *N. brasiliensis*-infected mice from the breedings described in (K). **(Q)** Frequencies of IL-13^+^, IFN-γ^+^ and IL-17A^+^ CD4^+^ T cells in the lungs of *N. brasiliensis*-infected mice from the breedings described in (K). **(B-J)** Data pooled from three independent litters per genotype; n= 19-20 mice per group. **(L-P)** Data pooled from three independent litters per maternal genotype; n= 12-14 mice per group. **(B-D; F; H; L-P)** Each symbol represents an individual mouse. Error bars represent Mean ±SD. **(J; Q)** Box-and-whisker plots display first and third quartiles, and the median; whiskers are from each quartile to the minimum or maximum. Normality of the samples was assessed with D’Agostino Pearson normality test; statistical analysis was then performed using Student’s t test or Mann-Whitney test. * *p* < 0.05; ** *p* < 0.01; *** *p* < 0.001

Next, we sought to evaluate if SCFA supplementation pre-weaning would also be able to suppress the increased inflammatory response during helminth infection observed in the progeny of TCRδ^-/-^ dams **(Figures 5A-J)**. Therefore, we infected with *N. brasiliensis* SCFA-treated mice, born from TCRδ^-/-^ dams, together with vehicle-treated mice, born from TCRδ^+/-^ dams **(Figure 5K)**. Critically, upon SCFA administration the offspring of TCRδ^-/-^ dams showed no differences, when compared to controls, in the numbers of total leukocytes **(Figure 5L)**, neutrophils **(Figure 5M)**, and eosinophils **(Figure 5N)** infiltrating the lung upon infection. Furthermore, we also found the number of infection-induced PD-L2-expressing macrophages **(Figure 5O)** and DCs **(Figure 5P)** to be similar between mice born from TCRδ^-/-^ and TCRδ^+/-^ dams. Finally, after SCFA supplementation, the progeny of TCRδ^-/-^ dams presented similar frequencies of lung IL-13^+^, IL17^+^, and IFN-γ^+^ CD4^+^ T cells in response to *N. brasiliensis* infection, when compared to mice born from TCRδ^+/-^ dams **(Figure 5Q)**. Taken together, our data ascribe the phenotypes observed in the TCRδ^-/-^ progeny to decreased maternal transfer of SCFA-producing bacteria, which beyond increasing first-breath-induced type-2 inflammation, exacerbates inflammatory responses to helminth infection after weaning.

## Discussion

Non-heritable factors, including the microbiome, are perceived as major contributors to systems physiology and immune variation in healthy humans *(Brodin et al., 2015; Stappenbeck and Virgin, 2016)*. In mice, many changes in cellular and humoral immune responses have been formally demonstrated to be microbiota-dependent *(Moon et al., 2015; Olszak et al., 2012)*. While this makes the design of experiments aimed at evaluating gene-intrinsic effects on a given phenotype more complex, it can be resolved by the use of littermate controls *(Stappenbeck and Virgin, 2016)*. However, a literature review from almost 3 decades of γδ T cell research in infection and immunity shows that only around 15% of the studies using TCRδ^-/-^ mice have used littermate controls **(Figure S2)**, which may explain some of the incongruences reported over the years. In the present study, a puzzling finding regarding differences in lung ILC2 in TCRδ^-/-^ mice when compared to age-, sex- and genetic background-matched C57BL/6J controls, but not in TCRδ+/+ and TCRδ-/- littermates, led us to investigate the factor influencing lung type-2 immunity in our TCRδ^-/-^ colony; and, most importantly, to enquire its root in the absence of maternal γδ T cells. By employing a reciprocal breeding strategy, we ascribed the vertical control of offspring pulmonary type-2 immunity by maternal γδ T cells to the transfer of microbiota during birth and fostering. The bacterial communities transferred by TCRδ^-/-^ dams possessed a discrete functional capacity, causing their pups to have decreased availability of specific microbial-derived SCFA, which regulated, via a gut – lung axis, type-2 inflammation in early life.

The fact that mice from the TCRδ^-/-^ colony had a very different fecal microbiota, when compared to their counter-parts coming from the TCRδ^+/-^ colony, was striking: first, because the heterozygous breeding was established with founder animals from the TCRδ^-/-^ colony; and second, because both mouse lines were kept under the same selective environmental pressures. Interestingly, a different study conducted in an unrelated vivarium reported significant differences in the oral microbiota of TCRδ^-/-^ mice when compared to genetic background-matched C57BL/6J mice *(Krishnan et al., 2018)*. Moreover, using a model for acute depletion of γδ T cells, another group found that continuous ablation of γδ T cells induces profound changes in the oral microbiome, compared to control (non-depleted) animals *(Wilharm et al., 2019)*. Altogether, these data suggest that γδ T cells – found in abundant numbers at barrier tissues – actively select bacterial commensals. Although underlying mechanisms are still largely unexplored *(Papotto et al., 2021)*, such γδ T cell-mediated microbiome selection could explain our findings that even after co-housing, TCRδ^-/-^ dams retain differences in cutaneous bacterial communities. This is in line with evidence that co-housing does not equalize niche-specific microbiota within the intestines of genetically identical mice *(Robertson et al., 2019)*. However, to properly address this important question, one requires a specific experimental design that takes into account the long-term evolution and short-term adaptation of the microbiota within a given host *(Yilmaz et al., 2021)*.

In what concerns maternal-offspring interactions, our results suggest that discrete bacterial communities present in the skin of TCRδ^+/-^ dams are vertically transferred during birth and nursing, promoting a SCFA-rich microbiome. Previous analysis of human mother-infant pairs showed that maternal skin and vaginal strains provide the earlier colonization, followed by gut-derived strains *(Ferretti et al., 2018)*. As γδ T cells are found in relatively high numbers in the female reproductive tract, it was somewhat surprising to not see differences in the vaginal microbiota of TCRδ^-/-^ and TCRδ^+/-^ dams. However, a caveat is that the vaginal wash used for the 16S rRNA-sequencing in the present study was not performed in pregnant females just before birth. Maternal vaginal microbiota changes significantly during gestation, with a particular dramatic change during the late phases of pregnancy and at birth *(Rasmussen et al., 2020)*. Further, even though maternal vaginal microbiota apparently has only modest influences on neonatal colonization *(Rasmussen et al., 2020)*, disruption of this process by cesarean-section, for instance, has shown to significantly change microbial colonization in newborns *(Backhed et al., 2015; Bokulich et al., 2016; Roswall et al., 2021; Tanaka and Nakayama, 2017)*. Of note, despite lung microbiota being shown to keep local type-2 immunity in check during allergen challenge *(Gollwitzer et al., 2014)*, we did not observe differences in the pulmonary microbiota of pups born from TCRδ^-/-^ and TCRδ^+/-^ dams, and our results thus support a skin(dam)-gut-lung(pup) axis regulating neonatal pulmonary inflammation.

Even though rodent pulmonary immune system development starts during fetal life *(McCarthy et al., 1992)*, the main changes occur during the first two weeks of life. Changing from a fluid-filled to an air-filled environment entails sustained basal type-2 immune responses – a critical feature to keep organ function during this heavy remodeling phase *(de Kleer et al., 2016; Saluzzo et al., 2017)*. The activation of a perinatal IL-33 – ILC2 – IL-13 axis due to mechanical damage caused by the first-breath reaction critically regulates resident populations such as alveolar macrophages *(Saluzzo et al., 2017)* and DCs *(de Kleer et al., 2016)*, with long-lasting effects in lung immunity. Our results indicate that early-life gut-colonizing microbes and their metabolites are able to fine-tune this lung type-2 inflammation, indicating that the process of IL-33 release by airway epithelial cells (AEC) involves more than physical stress, thus adding to the state-of-the art *(Saluzzo et al., 2017)*. In our model, the cells responding to the SCFA remain to be elucidated. Nonetheless, administration of the SCFA cocktail decreased both lung IL-33 levels and ILC2-derived IL-13, suggesting direct action on AEC, with downstream effects on ILC2 activation. Of note, production of cytokines by cell lines and primary cultured human AEC is modulated by SCFA in vitro *(Ghorbani et al., 2015)*. Moreover, butyrate, but not acetate, suppresses IL-13 production by ILC2 *(Thio et al., 2018)*. Although not much is known about the effects of pentanoate and hexanoate on immune responses, acetate can decrease type-2 inflammation in a mouse model of allergic asthma *(Trompette et al., 2014)*. In addition, feeding mothers with acetate, or with a high-fiber diet, which induces an acetate-producing microbiota, prevents the development of allergic asthma in the progeny by maternal transfer of this SCFA during pregnancy *(Thorburn et al., 2015)*. In our study however, in utero transfer of SCFA was not responsible for the regulation of lung type-2 immunity in the offspring. Finally, it is important to note that, whereas for the intestinal immune system the action of SCFA seem to occur at the time of weaning *(Al Nabhani et al., 2019)*, we show that lung immunity is affected by microbiota-derived SCFA before weaning, highlighting the discrete time maturation windows for the immune cells residing within different tissues.

While nutrition, especially during pregnancy and breast feeding, has a major impact on early microbiota composition and offspring physiology *(Backhed et al., 2015; Kimura et al., 2020; Thorburn et al., 2015)*, our study highlights the importance of microbiota transmission from mother-to-infant at the perinatal stage *(Ferretti et al., 2018)* to the establishment of gut microbiota functionality, which conditions the physiology of a distant organ, the lung, undergoing major immune remodeling during early life.

## Matherial and Methods

### Mice and breeding setups

C57BL/6J (B6) mice were purchased from Charles River Laboratories, C57BL/6J.TCRδ^-/-^ (referred to as TCRδ^-/-^, hereon in) were purchased from The Jackson Laboratory. C57BL/6J.JHT (referred as JHT) mice were obtained from Instituto Gulbenkian de Ciência (Oeiras, Portugal). Mice were bred and maintained in the specific pathogen-free animal facilities of Instituto de Medicina Molecular João Lobo Antunes (Lisbon, Portugal). After reaching sexual maturity (at around 8 weeks) TCRδ^-/-^ females were crossed with TCRδ^+/-^ males (obtained from the cross of TCRδ heterozygous animals derived, in turn, from a TCRδ^-/-^ x B6 breeding), and vice-versa; the pups obtained from the aforementioned crossings were kept with the parents until weaning, to be later divided by sex. All the pups were used at PN16±2, unless stated otherwise. All experiments involving animals were done in compliance with the relevant laws and institutional guidelines and were approved by local and European ethic committees.

### Cross-fostering and C-Sec surgeries

For cross-fostering experiments, mice were time-mated. Briefly, female mice were put in a cage containing the bedding of a cage containing male animals for a period of 48h. After this period of time, 1 or 2 females were introduced to a single-housed male. The following day, the presence of vaginal plugs (considered as E0.5) was recorded and males were transferred into a new cage. In some cases, the animals were left together for 2 days. Subsequently, pups born from a TCRδ^-/-^ dam were swapped with those from a TCRδ+/- dam at day 1 post-birth (and vice-versa), respecting litter sizes (differences of 3 pups maximum). For C-Sec experiments TCRδ^+/-^ and TCRδ^-/-^ females were bred with males from the opposing genotype in two groups, with an interval of 1 day between them. The group of females giving birth first was then defined as the surrogate mothers. Between days 18.5 and 19.5 the females from the second group underwent C-Sec surgery. Briefly, females are euthanized; the uterus is removed and submerged in iodine. Pups are taken out from the uterus on top of a sterile heated plate, cleaned and reanimated. When presenting regular respiratory movements, pups are put together with surrogate dams (from the same genotype of their own dam) and their progeny.

### Monoclonal antibodies

The following anti-mouse fluorescently labeled monoclonal antibodies (mAbs) were used (antigens and clones): CD3 (145.2C11, 17A2), TCRδ (GL3), CD90.2 (Thy1.2; 53-2.1), CD19 (6D5), CD23 (B3B4), CD45 (30F11), CD4 (GK1.5) NK1.1 (PK136), PD-L1 (10F.9G2), PD-L2 (TY25), CD11b (M1/70), ST2 (IL-33R; DIH9), CD11c (N418), Sca-1 (D7), Gr-1 (RB6-8C5), Ly6G (1A8), Ly6C (HK1.4), IA/IE (MHC II; M5/114.15.2), CD24 (M1/69), Siglec-F (E50-2440), CD64 (X54-5/7.1), CD206 (CO68C2), IL-13 (eBio13A), IL-9 (RM9A4), IL-5 (TRFK5). Lineage antibody cocktail (Lin) comprises a combination of the antibodies: CD3, CD4, CD19, CD23, NK1.1, CD11c and Gr-1. Antibodies were purchased from BD Biosciences, eBiosciences or BioLegend.

### Cell preparation, flow cytometry, and analysis

For cell surface staining, single-cell suspensions were incubated in presence of anti-CD16/CD32 (eBioscience) with saturating concentrations of combinations of the mAbs listed above. Lungs were dissected and tissue was cut into pieces, then digested with collagenase D (0.66 mg/ml; Roche) and DNase I (0.10 mg/ml) (Sigma-Aldrich) in RPMI 1640 containing 5% Fetal Bovine Serum (FBS) at 37°C for 30 min. For lamina propria cell preparation, small intestines were dissected, washed in ice cold PBS and Peyer’s patches were excised. The organ was then cut into pieces and incubated with EDTA 0.05M at 37°C for 20 min; cells were then washed and passed through a 100-µm cell strainer and then digested as the lungs. Single cells were isolated by passing the tissue through a 40-µm cell strainer, followed by a 70% / 20% Percoll (Sigma-Aldrich) gradient and 30-min centrifugation at 2400 rpm. Leukocytes were recovered from the interface, resuspended, and used for further analyses. For intracellular cytokine staining, cells were stimulated with PMA (phorbol 12-myristate 13-acetate) (50ng/mL) and ionomycin (1µg/mL), in the presence of Brefeldin A (10µg/mL) (all from Sigma) for 3h to 4h at 37°C. Cells were stained for the identified above cell surface markers, fixed 30min at 4°C and permeabilized with the Foxp3/Transcription Factor Staining Buffer set (eBioscience) in the presence of anti-CD16/CD32 (eBioscience) for 10min at 4°C, and finally incubated for 1h at room temperature with identified above cytokine-specific Abs in permeabilization buffer. Cells were analyzed using FACSFortessa (BD Biosciences) and FlowJo software (Tree Star). For cell sorting of lung interstitial macrophages (IM), lung cell suspensions were prepared and immuno-stained as described above. IM were defined as CD45^+^CD64^+^CD24^-^CD11b^+^CD11c^-^ live cells, and then electronically sorted on a FACSAria (BD Biosciences) into sterile RNase-free eppendorfs containing 200µL of lysis buffer (Qiagen), for posterior mRNA isolation.

### Immuno-detection of cytokines in lung tissue samples

Lung tissue was homogenized in 2 mL sterile tubes containing PBS plus a cocktail of protease inhibitors (cOmplete ULTRA tablets, Mini; Roche) and 1mm of diameter zirconia/silica beads (BioSpec Products), with the help of a tissue homogenizer. The homogenate was centrifuged in order to remove cellular debris and beads, and stored at -80°C until usage. Cytokine concentrations were assessed using LEGENDplex™ Mouse Th1/Th2 and Mouse Cytokine Panel 2 kits – both bead-based assays that use the principles of sandwich ELISA to quantify soluble analytes using a flow cytometer (Biolegend) – according to manufacturer’s instruction. Cytokine concentrations were normalized using total protein concentration in tissue homogenate, determined in a NanoDrop spectrophotometer (ThermoFisher).

### Immuno-detection of antibodies in serum samples

Blood was collected from the cheek pouch in adults, and from decapitation in pups. Serum samples were then stored at -20°C until usage. Determination of immunoglobulins isotypes and serum concentrations were assessed using LEGENDplex™ Mouse Immunoglobulin Isotyping Panel (Biolegend) according to manufacturer’s instruction.

### RNA isolation, cDNA preparation and real-time PCR

mRNA was prepared from tissue homogenates using RNeasy® Mini Kit (Qiagen). Reverse transcription was performed with random oligonucleotides (Invitrogen) using Moloney murine leukemia virus reverse transcriptase (Promega) for 1 hour at 42 °C. Relative quantification of specific cDNA species to endogenous reference *Hprt, Actb* or β*2microglobulin* was carried out using SYBR on ABI ViiA7 cycler (Applied Biosystems). The C_T_ for the target gene was subtracted from the average C_T_ for endogenous references, and the relative amount was calculated as 2^-CT^.

### N. brasiliensis infection

*N. brasiliensis* was maintained by serial passage through Sprague-Dawley rats, as previously described (Lawrence et al., 1996). Third-stage larvae (L3) were washed ten times with sterile PBS prior to subcutaneous infection of 300 L3s per mouse. On day 6 post-infection (p.i.) lungs were excised and processed accordingly to the analysis to be conducted.

### Antibiotic treatment and SCFA supplementation

TCRδ^-/-^ and TCRδ^+/-^ female and male adult mice were maintained with a mixture of antibiotics (5mg/ml of streptomycin, 1mg/ml of ampicilin, 1mg/ml of collistin, and 0.5mg/ml of vancomycin; Sigma Aldrich) in the drinking water for at least three weeks prior the start of the breedings, and then crossed as already described. Antibiotic treatment was kept for the whole duration of pregnancy and fostering. For direct supplementation of SCFA, 50µL of a cocktail containing acetate (2mg/mL), pentanoate (0.4mg/mL) and hexanoate (0.35mg/mL) was administered i.p. 3 times/week, starting at PN7 and finishing at PN13 for experiments analyzing the steady-state lung; and starting at PN14 and finishing at PN20 for experiments analyzing the response to *N. brasiliensis* infection. In both cases controls were injected with PBS. When given to dams, a SCFA cocktail containing acetate (50mM), pentanoate (37.5mM) and hexanoate (12.5mM) was added to the drinking water at day 2±1 after the litter was born and kept during the fostering period.

### Microbiota profiling via 16S rRNA gene sequencing

Samples were collected in sterilized surfaces using autoclaved materials. Fecal microbiota was assessed from two to three fecal pellets per animal. Skin microbiota was assessed upon removal of one ear per animal. Vaginal microbiota was assessed from the vaginal wash with 30µL of sterile PBS. Lung and cecum mucosa associated microbiota were assessed from the whole excised organs. Immediately after collection, samples were snap-frozen in liquid nitrogen and stored at -80°C until microbial DNA extraction. Before extraction, samples were homogenized in 2mL sterile tubes containing 1mm of diameter autoclaved zirconia/silica beads (BioSpec Products), with the help of tissue homogenizer. Microbial genomic DNA was extracted using the QIAamp® Fast DNA Stool Mini kit (Qiagen). The 16S rRNA V4 amplicons and ITS1-spanning amplicons were both generated using the following Earth Microbiome Project benchmarked protocols. Amplicons were then sequenced using on a 2×250bp PE mode Illumina MiSeq platform at Instituto Gulbenkian de Ciências (IGC) Genomics Unit (Oeiras, Portugal). Raw sequences obtained from 2 independent Illumina MiSeq runs were initially pooled, and analyzed in the QIIME pipeline version 2 using custom analysis scripts for analysis on the UBELIX Linux cluster of the University of Bern (High Performance Computing, University of Bern, Bern, Switzerland). Amplicon sequencing variants were assigned using Greengenes (13_8) databases using the feature-classifier classify-sklearn function with a 97% sequence identity threshold *(McDonald et al., 2012)*. The taxonomy, rep-seqs, and rooted-tree files generated in QIIME2 were called out in phyloseq pipeline in R. Calculation and plotting of the alpha diversity (Simpson and Shannon index), beta diversity (Bray-Curtis dissimilarities on NMDS plot), and statistical analysis using Adonis test were performed using phyloseq pipeline *(Bolyen et al., 2019; McMurdie and Holmes, 2013; Yilmaz et al., 2019)*. The multivariate analysis by linear models (MaAsLin2) was used to find associations between metadata and microbial community abundance (https://huttenhower.sph.harvard.edu/maaslin/) *(Mallick et al., 2021)*. Taxa that were present in ≥30% of the samples and had >0.001% of total abundance were set as the cut-off values for further analysis. After Benjamini-Hochberg false discovery rate correction, adjusted p-value (<0.2) was considered significant. The prediction of metagenome functional content was performed using 16S sequencing data in the Phylogenetic Investigation of Communities by Reconstruction of Unobserved States (PICRUSt) pipeline *(Langille et al., 2013)*. PICRUSt predictions were categorized as levels 1–3 and further into Kyoto Encyclopedia of Genes and Genomes pathways.

### SCFA measurement

Sample analysis was carried out by MS-Omics (www.msomics.com) as follows. Samples extracted by adding ultrapure water to the cecum (100µL to samples with less than 30mg and 150µL to samples with more). The filtered cecum water was further extracted by addition of deuterium labelled internal standards and mixing with MTBE directly in the injection vial. All samples were analyzed in a randomized order. Analysis was performed using a high polarity column (ZebronTM ZB-FFAP, GC Cap. Column 30m x 0.25mm x 0.25µm) installed in a GC (7890B, Agilent) coupled with a quadropole detector (5977B, Agilent). The system was controlled by ChemStation (Agilent). Raw data was converted to netCDF format using Chemstation (Agilent), before the data was imported and processed. The raw GC-MS data was processed by software developed by MS-Omics and collaborators. The software uses the powerful PARAFAC2 model and can extract more compounds and cleaner MS spectra than most other GC-MS software.

### Literature review

A search was performed in the PubMed data base in May/2019, using the keywords “gamma delta T cells AND infection” and “gamma-delta T cells AND infection”. Studies employing TCRδ^-/-^ mice were then divided regarding the use of littermates or genetic background-matched mice as controls. Journals were divided by their impact factor at the moment of the literature search.

### Statistical Analysis

In most experiments 3 independent litters were pooled together, unless stated otherwise. D’Agostino-Pearson normality test was applied to assess the normality of the populations analyzed; statistical significance of differences between populations was, then, assessed with the Student’s t-test or by using a two-tailed nonparametric Mann-Whitney U test, when applicable. For grouped statistical analysis, Two-Way ANOVA was employed. The *p* values <0.05 were considered significant and are indicated on the figures.

## ACKNOWLEDGEMENTS

We thank the precious technical assistance of the staff of iMM’s Rodent Facility – especially Iolanda Moreira, Pedro Santos and Bruno Novais – and Flow Cytometry Unit. We are also grateful to the Rodent Facility and Genomics Unit of Instituto Gulbenkian de Ciência, and MS-Omics for the services provided. We thank Matthew R. Hepworth and Marc Veldhoen for critical reading of the manuscript; and Debanjan Mukherjee, Helena Brigas, Julie C. Ribot, Karine Serre, Ana Pamplona, Julie Darrigues (iMM), Rita Domingues, Julie Chesné, Vânia Cardoso, Henrique Veiga-Fernandes (Champalimaud Center for the Unknown), James Parkinson (University of Manchester) and Ziad Al Nabhani (University of Bern) for helpful discussions and technical advice. This work was funded by Fundação para a Ciência e Tecnologia (PTDC/MED-ONC/6829/2020 to B.S.-S.; PD/BD/105855/2014 to P.H.P.), European Molecular Biology Organization (LTF 191-2019 to P.H.P and ALTF 252-2017 to G.J.F); European Commission Marie Sklodowska-Curie Individual Fellow-ship (ref. 752932 to G.J.F); Swiss National Foundation (SNF) Ambizione Grant (PZ00P3_185880 to B.Y.), Novartis Foundation for Medical-Biological Research (#19A013 to B.Y); and the Wellcome Trust (106898/A/15/Z to J.E.A.).

## AUTHOR CONTRIBUTIONS

Conceptualization: P.H.P. and B.S.-S.; Methodology and Investigation: B.Y., G.P., S.M., C.C., G.F., D.G.C., N.G.-S., B.H.K.C., B.B., T.C., B.H.K.C., A.J.M., and J.E.A.; Data Analysis: P.H.P., B.Y. and G.P.; Resources: B.H.K.C. and J.E.A.; Writing – original draft: P.H.P. and B.S.-S.; Writing – review and editing: P.H.P., B.Y., J.E.A., and B.S.-S.

## Supplementary Figures

**Fig. S1.**
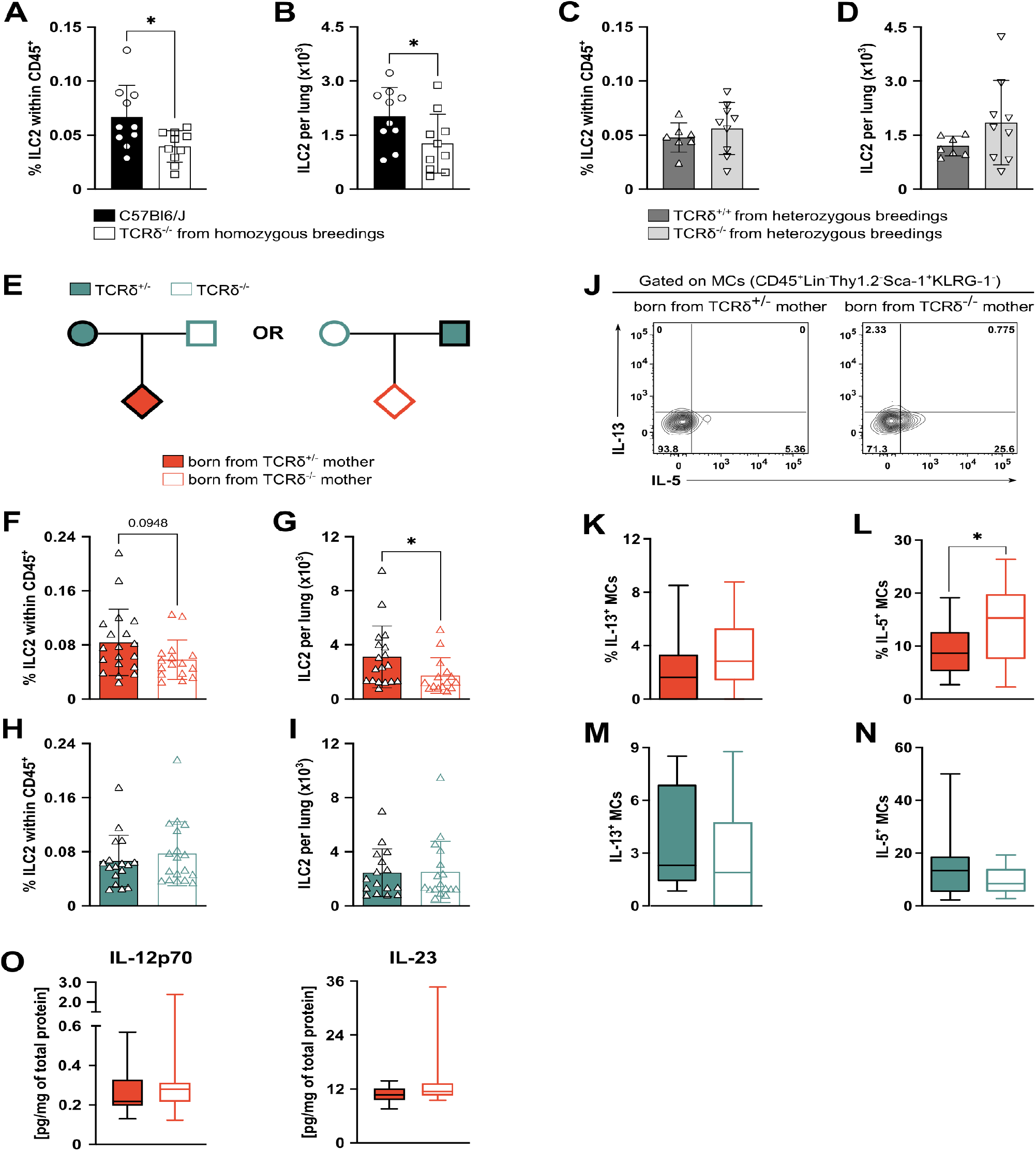
The offspring of TCRδ^-/-^ dams present differences in pulmonary type-2 immune system. **(A)** Frequencies and textbf(B) numbers of ILC2 (defined as Lin^-^Thy1.2^+^Sca-1^+^KLRG-1^+^ cells) isolated from the lungs of age- and sex-matched C57Bl/6J and TCRδ^-/-^. **(C)** Frequencies and textbf(D) numbers of ILC2 (defined as Lin^-^Thy1.2^+^Sca-1^+^KLRG-1^+^ cells) isolated from the lungs of TCRδ^+/+^ and TCRδ^-/-^ littermates from heterozygous breedings. **(E)** Frequencies and **(F)** numbers of ILC2 (defined as Lin^-^Thy1.2^+^Sca-1^+^KLRG-1^+^ cells) isolated from the lungs of pups from the breedings described in (Figure 1A), grouped by maternal genotype. **(G)** Frequencies and **(H)** numbers of ILC2 (defined as Lin^-^Thy1.2^+^Sca-1^+^KLRG-1^+^ cells) isolated from the lungs of pups from the breedings described in (Figure 1A), grouped by offspring genotype. **(I)** Flow cytometry analysis of intracellular IL-5 and IL-13 expression in mast cells (MCs; defined as Lin^-^Thy1.2^-^Sca-1^+^KLRG-1^-^ cells) isolated from the lungs of pups generated in the breedings described in (Figure 1A) and stimulated in vitro for 3h in the presence of PMA, Ionomycin and Brefeldin A. Frequencies of **(J)** IL-13^+^ and **(K)** IL-5^+^ cells within the MC population in the lungs of pups depicted in (Figure 1A), grouped by maternal genotype. Frequencies of **(L)** IL-13^+^ and (M) IL-5^+^ cells within the MCs in the lungs of pups depicted in (Figure 1A), grouped by offspring genotype. **(N)** Concentration of cytokines within the whole lung homogenate normalized by total protein in the tissue of pups from the breedings described in (Figure 1A), grouped by offspring genotype. (A-B) Data pooled from three independent experiments; n= 10 mice per group. **(C-D)** Data pooled from three independent experiments; n= 7-9 mice per group. **(E-M)** Data pooled from three independent litters per maternal genotype; n= 16-17 mice per group. (N) Data pooled from at least three independent litters per genotype; n= 18-23 mice per group. **(A-H)** Each dot represents an individual mouse; error bars represent Mean ±SD. **(J-N)** Box-and-whisker plots display first and third quartiles, and the median; whiskers are from each quartile to the minimum or maximum. Normality of the samples was assessed with D’Agostino Pearson normality test; statistical analysis was then performed using Student’s t test or Mann-Whitney test. * *p* < 0.05

**Fig. S2.**
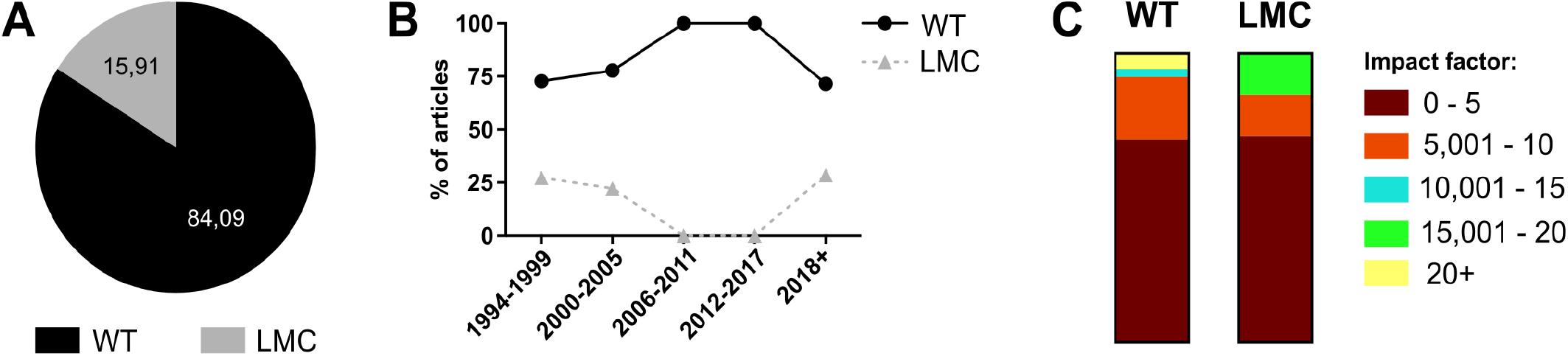
Littermate control usage in publications on γδ T cells in infection immunity. **(A)** Percentage of the usage of littermate versus genetic background-matched (“wild type”, WT) controls in studies employing TCRδ^-/-^ mice in experimental models of infection. Breakdown of the results presented in **(A)** over time **(B)** or by impact factor of the journal where the study was published **(C)**.

**Fig. S3.**
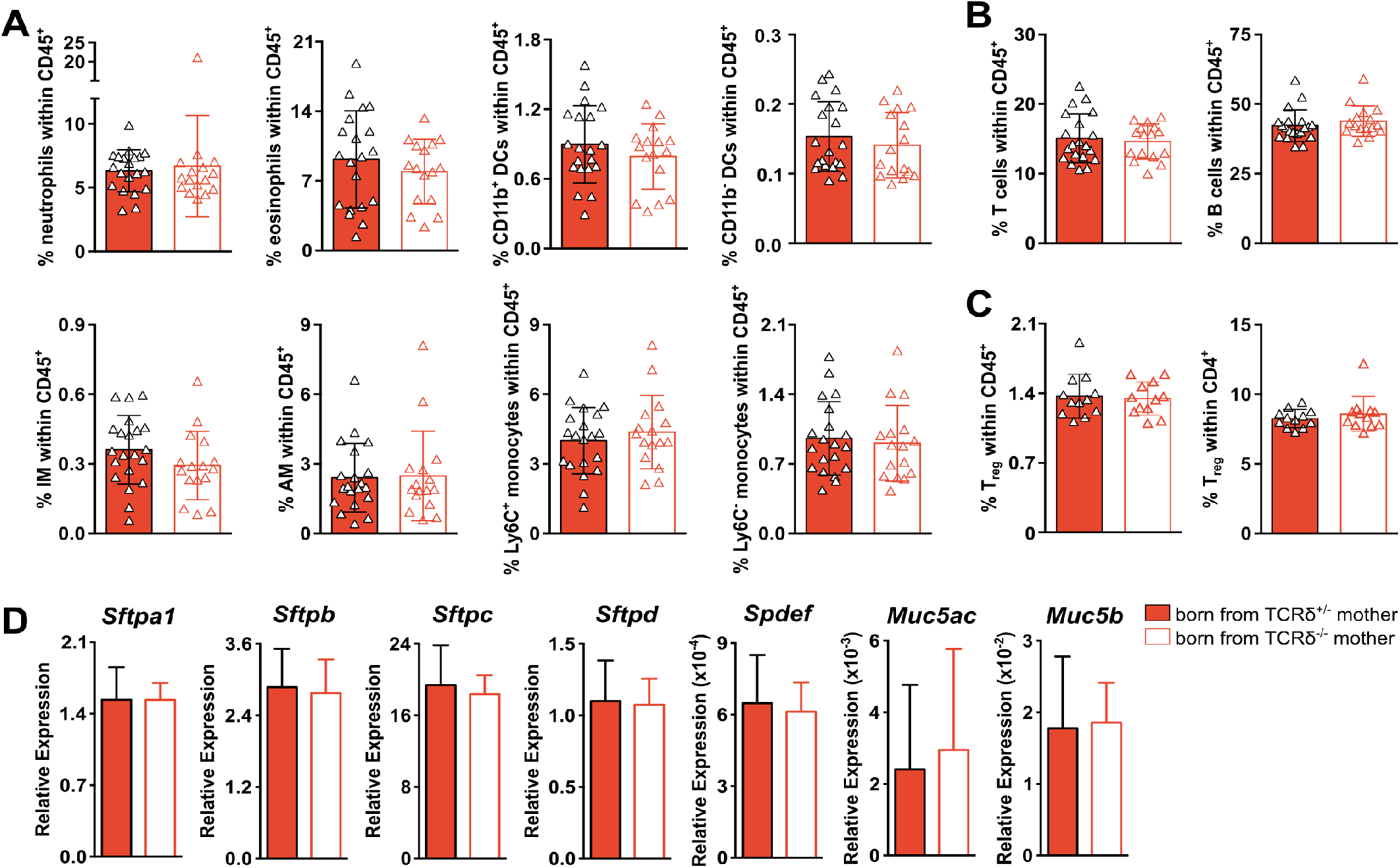
Absence of maternal γδ T cells does not change the distribution of immune cell subsets in the neonatal lung, nor the expression of epithelial genes. **(A)** Frequencies of lung innate immune cell populations within total CD45^+^ cells in pups obtained from the breedings depicted in (Figure 1A), grouped by maternal genotype. **(B)** Frequencies of lung T and B lymphocyte populations within total CD45^+^ cells in pups obtained from the breedings depicted in (Figure 1A), grouped by maternal genotype. **(C)** Frequencies of lung CD3^+^CD4^+^Foxp3^+^ Treg cells within total CD45^+^ cells (left panel) and within total CD4^+^ cells (right panel) in pups obtained from the breedings depicted in (Figure 1A). **(D)** Gene expression of epithelial genes in total lung tissue from in pups obtained from the breedings depicted in (Figure 1A) normalized to *Hprt*. Absolute numbers of lung-resident populations in pups obtained from the breedings depicted in (Figure 1A) **(A-B)** Data pooled from three independent litters per maternal genotype; n= 20-16 mice per group. **(C)** Data pooled from two independent litters per maternal genotype; n= 12 mice per group. **(D)** Data representative of two independent litters per maternal genotype, n= 11-9 mice per group **(A-C)** Each symbol represents an individual mouse. Error bars represent Mean ±SD. Normality of the samples was assessed with D’Agostino Pearson normality test; statistical analysis was then performed using Student’s t test or Mann-Whitney test.

**Fig. S4.**
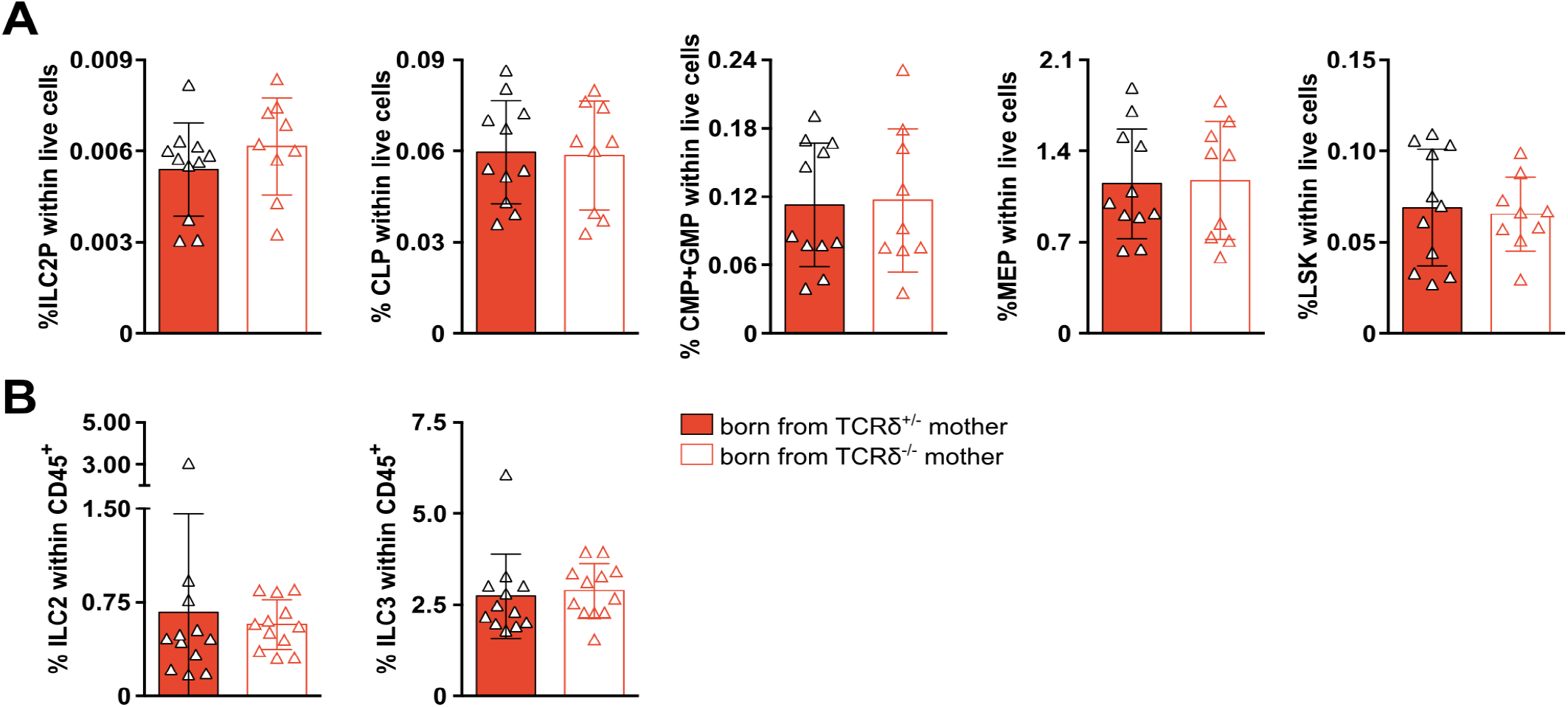
Absence of maternal γδ T cells does not change ILC populations in the small intestine, nor immune cell precursors in the bone marrow. **(A)** Frequencies of immune cell precursors (ILC2 precursors, ILC2P; common lymphoid progenitors, CLP; common myeloid progenitors + granulocyte-macrophage progenitors, CMP+GMP; megakaryocyte-erythroid progenitors, MEP; Lineage^-^Sca-1^+^c-Kit^+^ cells, LSK) in the bone marrow of pups obtained from the breedings depicted in (Figure 1A), grouped by maternal genotype. **(B)** Frequencies within total CD45^+^ cells of ILC2 (left panel) and ILC3 (right panel) in the small intestine lamina propria in pups obtained from the breedings depicted in (Figure 1A), grouped by maternal genotype. **(A)** Data pooled from two independent litters per maternal genotype; n= 9-11 mice per group. **(B)** Data pooled from two independent litters per maternal genotype; n= 12 mice per group. **(A-B)** Each symbol represents an individual mouse; error bars represent Mean ±SD. Normality of the samples was assessed with D’Agostino Pearson normality test; statistical analysis was then performed using Student’s t test or Mann-Whitney test.

**Fig. S5.**
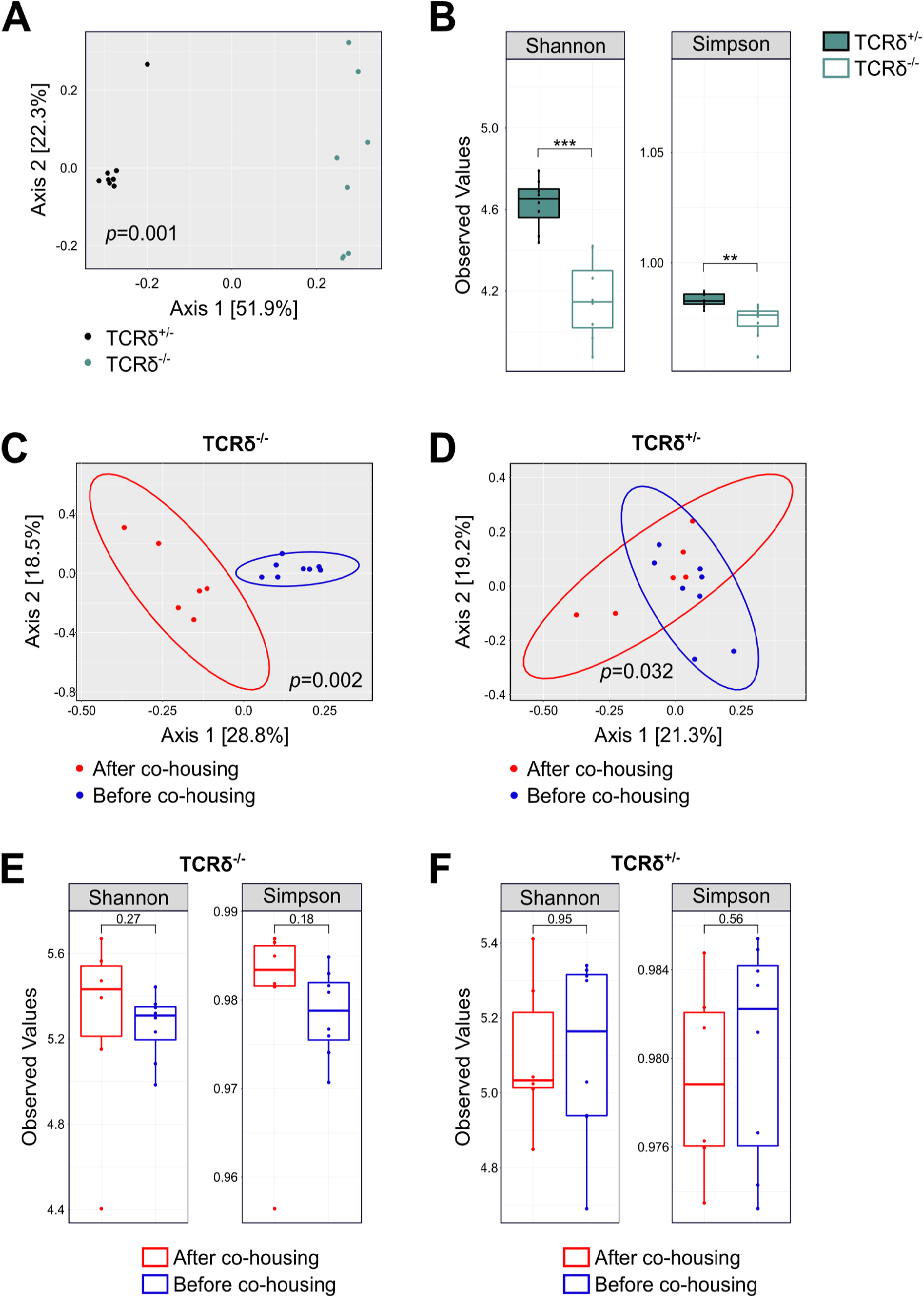
Co-housing of TCRδ^+/-^ and TCRδ^-/-^ dams with males from the opposing genotype changes their microbiota profile. **(A)** Microbial clustering is shown based on Bray–Curtis dissimilarity principal coordinate analysis (PCoA) beta diversity metrics using fecal samples from TCRδ^+/-^ and TCRδ^-/-^ female mice **(B)** Microbial composition differences in fecal microbiota from TCRδ^+/-^ and TCRδ^-/-^ female mice plotted using alpha diversity (Shannon and Simpson indices). Microbial composition differences in TCRδ^-/-^ **(C and E)** and TCRδ^+/-^ **(D and F)** dams before and after co-housing with males from the opposite genotype plotted using **(E-F)** alpha diversity (Shannon and Simpson indices) and **(C-D)** beta diversity (Bray–Curtis dissimilarity PCoA). Ellipsoids in PCoA plots represent a 95% confidence interval surrounding each group. **(A, C-D)** Non-parametric analysis of variance (Adonis) was used to test for significant difference between groups on PCoA plot. **(A-B)** n= 8 mice per group **(C-F)** n= 6-8 mice per group

**Fig. S6.**
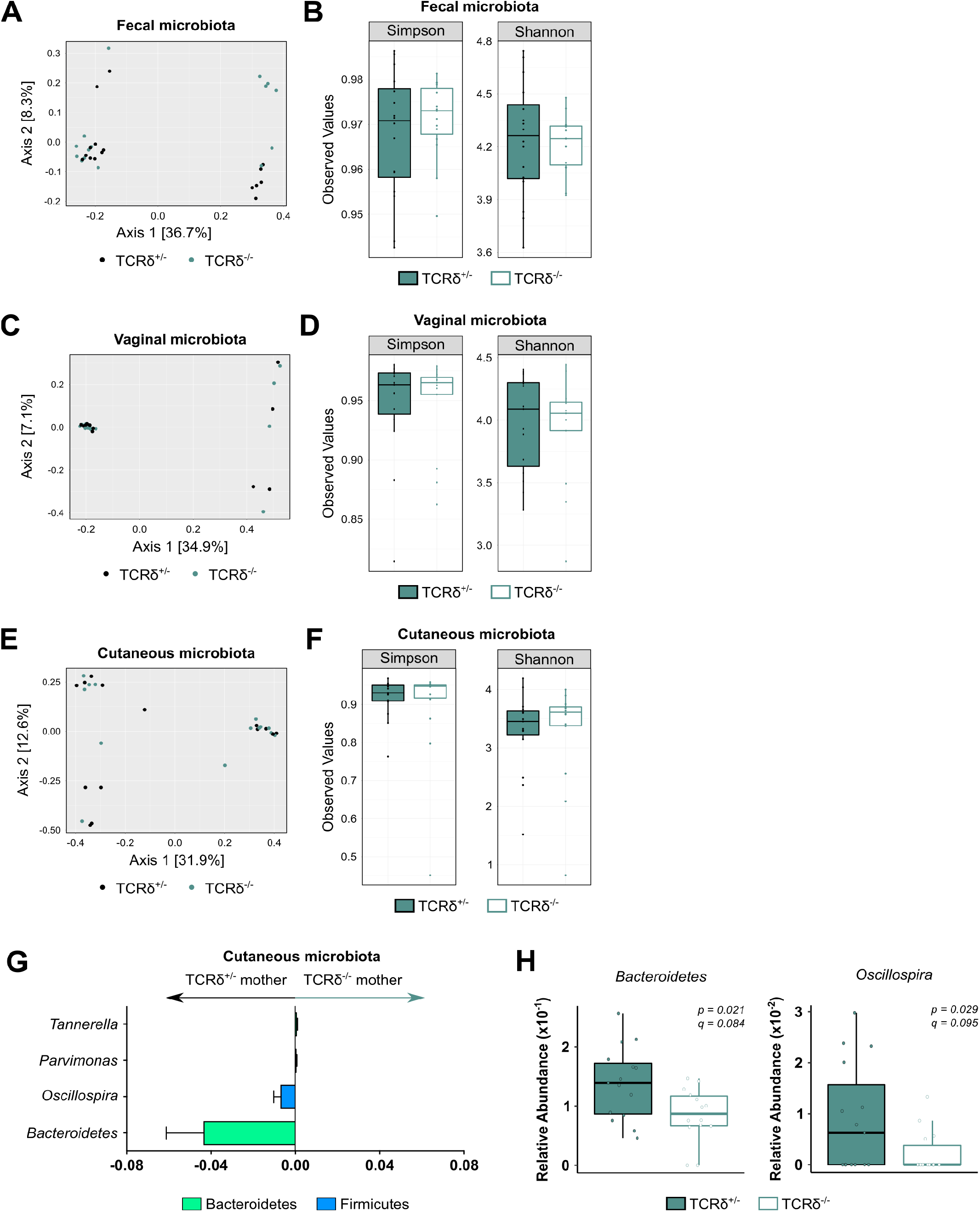
TCRδ^+/-^ and TCRδ^-/-^ dams display a discrete skin microbiota profile even after co-housing with males from the opposing genotype. Microbial composition differences in **(A-B)** fecal, **(C-D)** vaginal, and (E-F) skin microbiota from TCRδ^+/-^ and TCRδ^-/-^ dams are plotted using **(B, D and F)** alpha diversity (Shannon and Simpson indices) and **(A, C and E)** beta diversity (Bray–Curtis dissimilarity PCoA). **(G)** Skin microbiota taxa associated with TCRδ^+/-^ and TCRδ^-/-^ dams and plotted as relative abundance ratios. **(H)** Relative abundance of taxa significantly enriched (p < 0.05, q < 0.2) in the skin microbiota of TCRδ^+/-^ dams; Box-and-whisker plots display first and third quartiles, and whiskers are from each quartile to the minimum or maximum. **(A-H)** n= 14-15 mice per group.

**Fig. S7.**
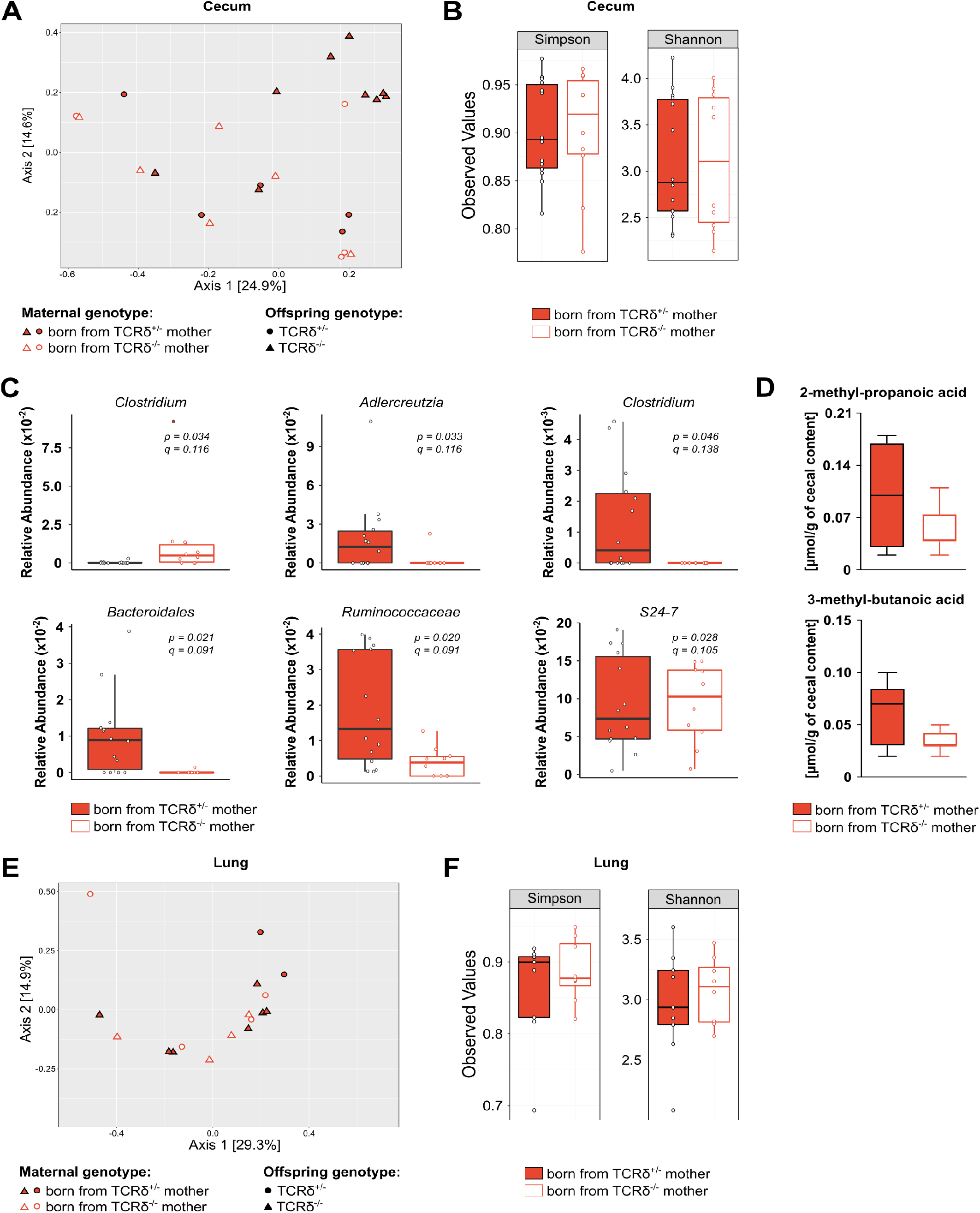
The offspring of TCRδ^+/-^ and TCRδ^-/-^ dams display discrete bacterial taxa composing their intestinal microbiota. Microbial composition differences in cecum microbiota from pups born from TCRδ^+/-^ and TCRδ^-/-^ dams are plotted using **(A)** beta diversity (Bray–Curtis dissimilarity PCoA) and **(B)** alpha diversity (Shannon and Simpson indices). **(C)** Relative abundance of taxa present in the cecum samples that were significantly different (p < 0.05, q < 0.2) between pups born from TCRδ^+/-^ and TCRδ^-/-^ dams. **(D)** Concentration of SCFA present in the cecum of pups born from TCRδ^+/-^ and TCRδ^-/-^ dams normalized by cecal content. Microbial composition differences in lung microbiota from pups born from TCRδ^+/-^ and TCRδ^-/-^ dams are plotted using **(E)** beta diversity (Bray–Curtis dissimilarity PCoA) and **(F)** alpha diversity (Shannon and Simpson indices). **(A-C)** Data pooled from at least two independent litters per maternal genotype; n= 10-14 mice per group. **(D)** Data from one litter per maternal genotype; n= 9-10 mice per group. **(E-F)** Data pooled from two independent litters per maternal genotype; n= 8-9 mice per group. **(B-D; F)** Box-and-whisker plots display first and third quartiles, and whiskers are from each quartile to the minimum or maximum.

**Fig. S8.**
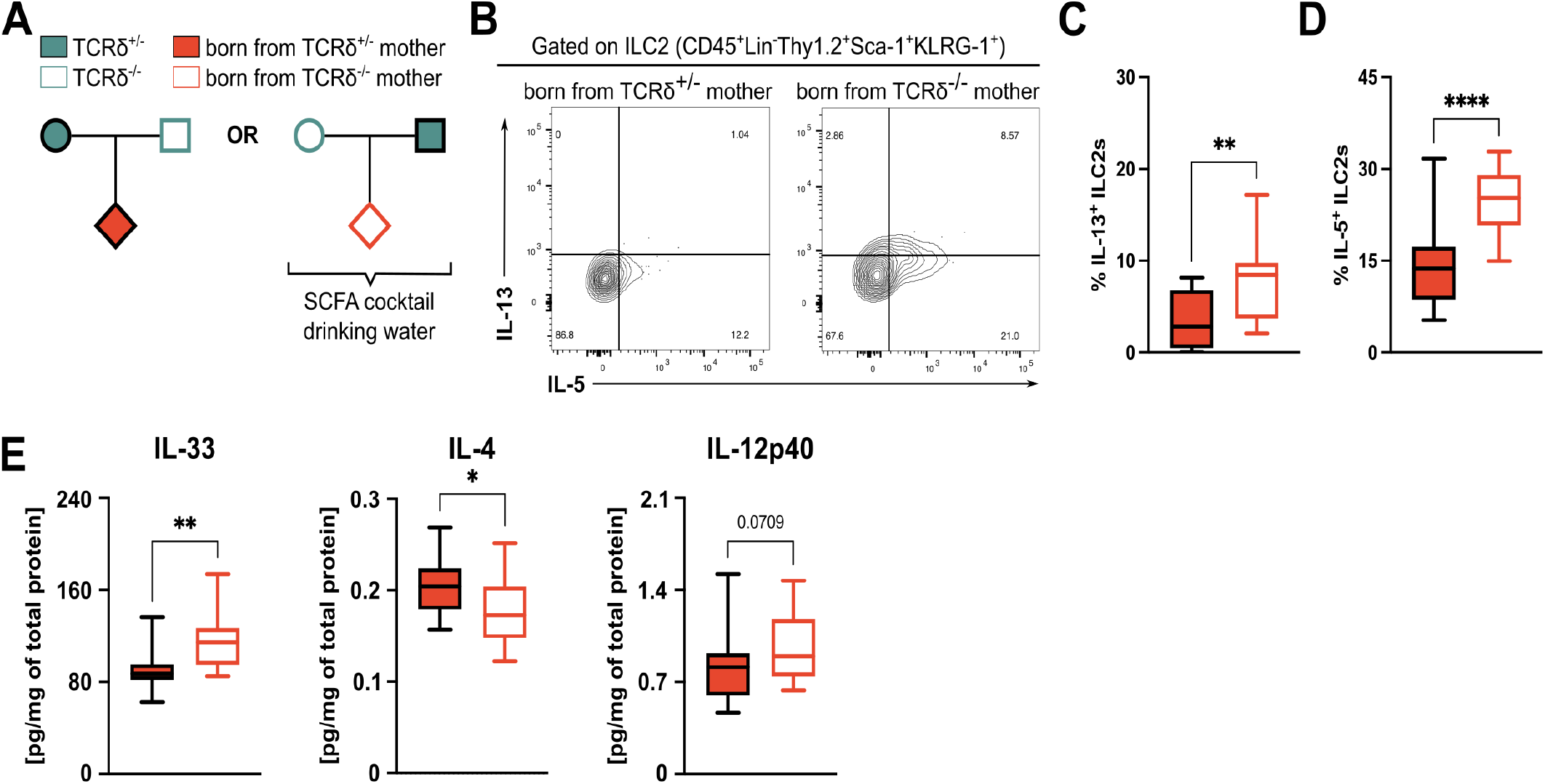
Maternal transfer of SCFA to the progeny does not completely abrogate perinatal lung type-2 inflammation. **(A)** Breeding strategy employed to evaluate the role of SCFA in regulating perinatal lung inflammation. TCRδ^+/-^ dams were crossed with TCRδ^-/-^ males, and vice-versa; pups were kept with the parents and a SCFA cocktail (acetate, pentanoate and hexanoate) was added to the drinking water immediately after birth. Analysis of neonatal lung immune system was conducted at PN16±2. **(B)** Flow cytometry analysis of intracellular IL-5 and IL-13 in ILC2s (defined as Lin^-^Thy1.2^+^Sca-1^+^KLRG-1^+^ cells) isolated from the lungs of pups generated in the breedings described in (A) and stimulated in vitro for 3h in the presence of PMA, Ionomycin and Brefeldin A. Frequencies of **(C)** IL-13^+^ and **(D)** IL-5^+^ cells within the ILC2 population in the lungs of pups depicted in (A), grouped by maternal genotype. **(E)** Concentration of cytokines within the whole lung homogenate normalized by total protein in the tissue of pups from the breedings described in (A), grouped by maternal genotype. **(B-E)** Data pooled from three independent litters per maternal genotype; n= 13-19 mice per group. **(C-E)** Box-and-whisker plots display first and third quartiles, and the median; whiskers are from each quartile to the minimum or maximum. Normality of the samples was assessed with D’Agostino Pearson normality test; statistical analysis was then performed using Student’s t test or Mann-Whitney test. * *p* < 0.05; ** *p* < 0.01; **** *p* < 0.0001

**Fig. S9.**
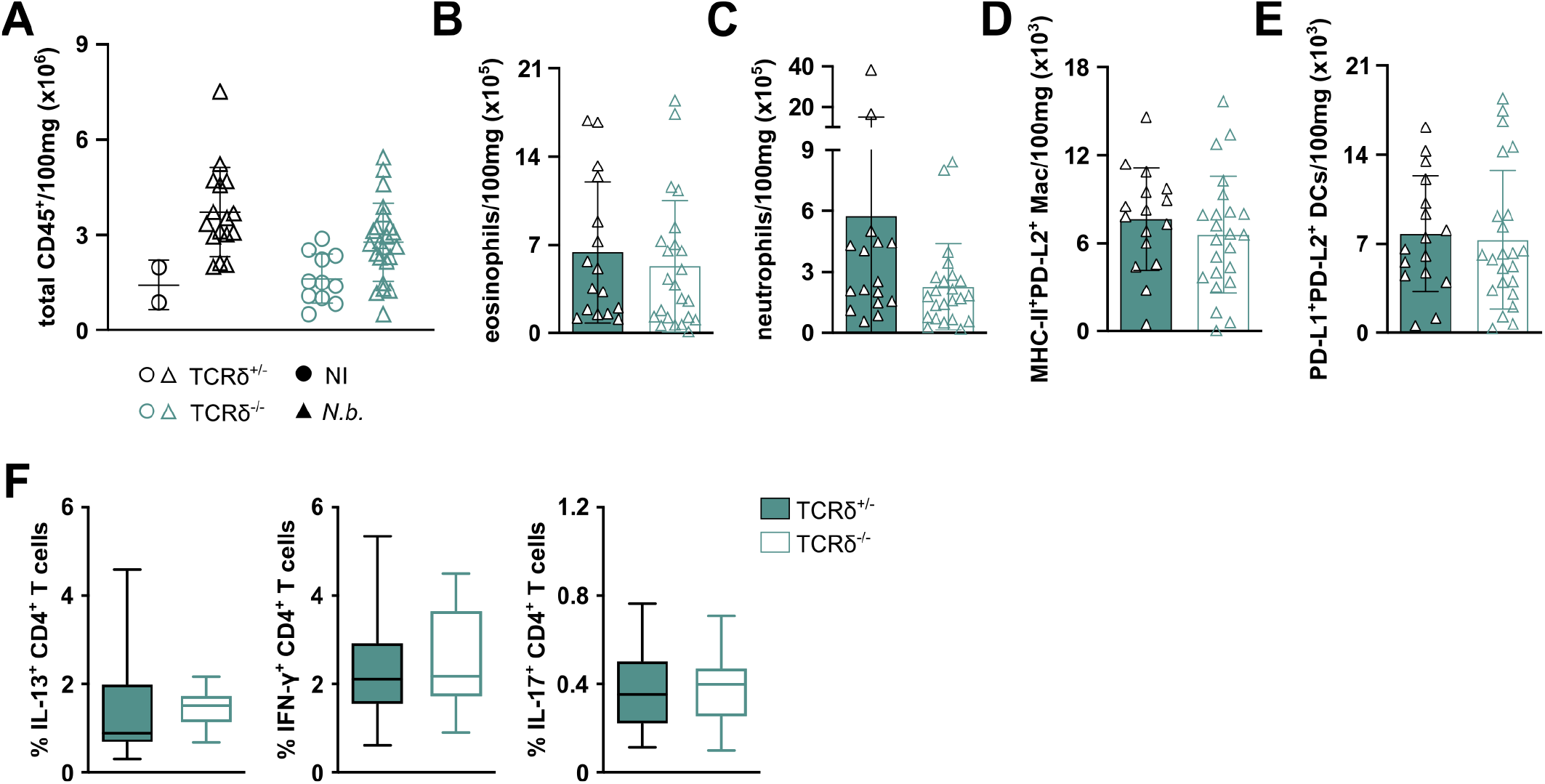
γδ T cell-deficient and -suficient mice respond similarly to N. brasiliensis infection. Numbers (normalized to 100mg of lung tissue) of **(A)** total leukocytes, **(B)** eosinophils, **(C)** neutrophils, **(D)** MHC-II^+^PD-L2^+^ macrophages and **(E)** PD-L1^+^PD-L2^+^ DCs in the lungs of *N. brasiliensis*-infected mice (Figure 5A). **(F)** Frequencies of IL-13^+^, IFN-γ^+^ and IL-17A^+^ CD4^+^ T cells in the lungs of *N. brasiliensis*-infected mice (Figure 5A). **(A-F)** Data pooled from three independent litters per genotype; n= 16-23 mice per group. Normality of the samples was assessed with D’Agostino Pearson normality test; statistical analysis was then performed using Student’s t test or Mann-Whitney test.

